# RNA 3’end tailing safeguards cells against products of pervasive transcription termination

**DOI:** 10.1101/2024.03.15.584960

**Authors:** Guifen Wu, Jérôme O. Rouvière, Manfred Schmid, Torben Heick Jensen

**Affiliations:** Department of Molecular Biology and Genetics, Universitetsbyen 81, Aarhus University, Aarhus, Denmark; QIAGEN Aarhus A/S, Silkeborgvej 2, Aarhus 8000, Denmark

**Keywords:** NEXT, RNA exosome, RNA U-tailing, RNA A-tailing, RNA decay

## Abstract

Premature transcription termination yields a wealth of unadenylated (pA^-^) RNA. Although this can be targeted for degradation by the Nuclear EXosome Targeting (NEXT) complex, possible back-up pathways remain poorly understood. Here, we find RNA 3’end uridylation and adenylation upon NEXT inactivation. U-tailed RNAs are generally short and modified by the cytoplasmic tailing enzymes, TUT4/7, following their PHAX-dependent nuclear export and prior to their degradation by the cytoplasmic exosome or the exoribonuclease DIS3L2. Longer RNAs are instead adenylated redundantly by enzymes TENT2, PAPOLA and PAPOLG. These transcripts are either degraded via the nuclear Poly(A) tail eXosome Targeting (PAXT) connection or exported by conventional factors and removed by the cytoplasmic exosome in a translation-dependent manner. Failure to do so decreases global translation and induces cell death. We conclude that post-transcriptional 3’end modification and removal of excess pA^-^ RNA is achieved by tailing enzymes and export factors shared with productive RNA pathways.

## Introduction

Mammalian genomes are pervasively transcribed by RNA polymerase II (RNAPII), yielding functional 5’end capped RNA along with a vast abundance of processing and transcriptional by-products ^1,2^. Given this wealth of RNA there is a high demand for efficient turnover of excess transcripts and various decay pathways have evolved to maintain RNA homeostasis by clearing cells of differently sized transcripts ^3,4^.

In the cytoplasm, the major pathways to control levels of translation-competent RNA involve 5’-3’ exonucleolysis, of prior de-capped RNA, by the XRN1 ribonuclease as well as 3’-5’ exonucleolysis by the RNA exosome, via its cytoplasm-specific exonuclease DIS3L ^5,6^ and its activating Superkiller (SKI) adaptor complex ^7–9^. RNAs undergoing such decay are most often 3’end processed and polyadenylated in the nucleus and therefore require cytoplasmic deadenylation as a necessary step in their turnover ^10,11^. Cytoplasmic RNA control can also occur by its translation-independent 3’end uridylation and subsequent 3’-5’ degradation by DIS3L2, which is independent on the RNA exosome ^12–14^. Here, RNA substrates range from certain mRNA to a variety of smaller noncoding RNA (ncRNA), which are uridylated by the uridylyl transferases TUT4 (ZCCHC11) and TUT7 (ZCCHC6) ^15–17^.

Prior to their cytoplasmic degradation, capped RNAs are exported from the nucleus through different pathways, depending on the RNA species and its features ^18,19^. Broadly speaking, the phosphorylated adaptor for RNA export (PHAX) is central for the export of short RNA, such as U-rich small nuclear RNA (UsnRNA), perhaps due to the affinity of PHAX for RNAs < 300 nt^20–23^. Longer RNAs, including mRNA and long ncRNA, are primarily exported in conjunction by the transcription-export (TREX) complex and the nuclear RNA export factor 1 (NXF1) ^24,25^.

In the nucleus, decay of capped transcripts is predominantly managed by the 3′-5′ exonucleolytic activity of the RNA exosome, harboring its nuclear-specific subunits DIS3 and RRP6 ^4,26^. Substrate access is facilitated by the exosome-associated RNA helicase MTR4 ^27–29^, which connects to one of two primary nucleoplasmic adaptors: the nuclear exosome targeting (NEXT) complex ^30^ or the polyA (pA) tail exosome targeting (PAXT) connection ^31,32^. NEXT forms a dimer of MTR4-ZCCHC8-RBM7 heterotrimers ^33,34^ and can target the exosome to short, TSS-proximal, pA^-^ transcripts via its connection to the 5’cap-binding complex (CBC) ^35–37^. Conversely, PAXT consists of a core MTR4-ZFC3H1 heterodimer, that associates with the nuclear pA binding protein (PABPN1) to facilitate the exosome-mediated decay of a variety of pA^+^ RNAs ^31,32,38–40^. By their combined targeting of pA^-^ and pA^+^ RNAs, NEXT and PAXT are believed to provide for much of the nuclear control of RNAPII-derived transcripts.

The above mentioned cytoplasmic and nuclear decay pathways have all been characterized in their own rights, but only little is known about the extent to which they functionally crosstalk and/or compensate for one another. This is, however, an important consideration as no pathway will, on its own, be fully sufficient. Moreover, changing gene expression patterns, occurring during cellular transition or upon stress, will unavoidably challenge a specific pathway with the potential hazard of overwhelming it. Finally, certain cellular conditions might affect RNA decay pathway components as exemplified by the NEXT subunit RBM7, which is phosphorylated upon DNA damage, leading to its decreased substrate targeting ^41–43^. Here, compensatory pathways are likely required to deal with accumulating NEXT substrates. In one example of NEXT pathway compensation, we previously showed that decreased NEXT activity may allow for the post-transcriptional polyadenylation of formerly pA^-^ NEXT substrates, leading to their PAXT-dependent nuclear turnover ^36^. Hence, this provides one possibility for fail-safe decay in case a first line of substrate recognition is prohibited. However, the cell may contain additional back up mechanisms for NEXT activity. This is because mammalian transcription units (TUs) broadly are prone to premature transcription termination, giving rise to shortened transcripts with either pA^+^ or pA^-^ 3’ends ^44–46^. Focusing here on pA^-^ RNAs, which are the targets of NEXT, early RNAPII termination within the first few kilobase (kb) of transcriptional progress is typically directed by the Integrator (INT)- or Restrictor (RES) complexes ^47–52^. Since both INT and RES operate genome-wide and in largely sequence-independent manners, extensive amounts of NEXT-substrates are constantly being produced.

Here, we explore which decay pathways maintain RNA homeostasis when NEXT-mediated turnover of pA^-^ RNA is compromised. Focusing on prematurely terminated and promoter upstream transcripts (PROMPTs), we show that the shorter fraction of these are exported, via PHAX, to the cytoplasm, where they are uridylated by TUT4/7 and degraded by the cytoplasmic exosome or the exoribonuclease DIS3L2. The fraction of relatively longer transcripts, escaping NEXT activity, are adenylated redundantly by the TENT2, PAPOLA and PAPOLG enzymes. This yields two possible fates; i) handover in the nucleus for PAXT-mediated decay, or ii) nuclear export by PHAX and the conventional mRNA export machinery and turnover by the cytoplasmic exosome.

## Results

### Short 3’end uridylated RNAs accumulate upon NEXT- and exosome-depletions

Inspired by our previous discovery that pA^-^ RNA substrates can undergo post-transcriptional polyadenylation in NEXT and exosome depletion conditions ^36^, we wondered whether other 3’end modifications might also occur. Hence, we scrutinized previous RNA 3’end sequencing (3’end seq) data ^36^, taking advantage of their *in vitro* polyadenylation by *E. coli* poly(A) polymerase (E-PAP), thus enabling the detection of non-canonical tails (Figure S1A). To obtain a general overview, we first combined data from four different siRNA knockdown conditions (siGFP, siRRP40 (also called siEXOSC3), siZCCHC8 and siZFC3H1) each containing biological triplicate steady-state (‘total’)- and duplicate 4-thiouridine (‘4sU’)-labeled RNA samples. From these data, non-aligned terminal sequences were collected and stratified by length, nucleotide content and the total number of reads computed. Filtering canonical pA tails from the analysis (see Methods), the interrogated RNAs contained a variety of non-templated tails, with ‘mixed A-tails’ of occasional non-adenosine residues residing inside pA tails (i.e A_n_G, A_n_U, A_n_C) and pure U-tails (U_n_) being the most abundant (Figure S1B). Although, one should be mindful about possible sequencing artefacts, similar observations were also made using different RNA sequencing techniques, such as TAIL-seq and 3’RACE-seq ^53,54^, with mixed A-tails and U-tails reported to be synthesized by non-canonical pA polymerases TENT4A/B ^53,55,56^ and terminal uridylyl transferases TUT4/7 ^57–59^, respectively

We decided to focus our efforts on U-tailing. To evaluate the extent to which U-tails might constitute sequencing artifacts, we first scored their detection frequencies in cellular RNA compared to that of the spike-in RNA lacking 3’end U-residues (Figure S1A). Cellular RNAs with 1-2 non-templated U’s were only marginally better detected than non-uridylated spike-in RNAs, while the difference became prominent for RNAs with ≧ 3 non-templated U’s (Figure S1C). The low detection-specificity for 1-2 U’s might relate to sequencing errors at read ends and/or other unknown technical issues ^60,61^. Regardless the cause, we subsequently focused only on RNAs with 3-9 residues long U-tails and measured their frequencies in response to depletion of the RNA exosome or its nucleoplasmic NEXT- and PAXT-adaptors. Notably, individual depletion of both the RNA exosome subunit RRP40 (siRRP40), and the NEXT component ZCCHC8 (siZCCHC8), led to increased cellular RNA U-tail frequencies of both total- and 4sU-RNA samples in an E-PAP-specific manner (Figure 1A). Conversely, depletion of the PAXT component ZFC3H1 (siZFC3H1) only affected 4sU RNA U-tail frequencies slightly. To directly visualize RNA U-tail abundance, we constructed aggregate plots, displaying their log_2_-transformed sequencing coverage (log_2_(cov)) when anchored to all annotated transcription start sites (TSSs). Similar to the frequency measurements, RNA U-tail abundance was increased upon exosome-and NEXT-depletions, but only mildly in 4sU RNA samples upon PAXT-depletion (Figure 1B). The fact that RNA U-tails were less upregulated in NEXT-*vs.* exosome-depletion conditions, when interrogating total RNAs, but reached similar levels in 4sU RNA samples, was reminiscent of our previous observation for other NEXT-sensitive targets, such as PROMPTs ^30,35,36^. This presumably reflects a redundancy between NEXT and PAXT, where NEXT-substrates can be polyadenylated and subjected to PAXT-mediated exosome decay ^36^. The similar observation for U-tailed RNA indicated multiple layers of RNA quality control, which we decided to address later.

**Figure 1.**
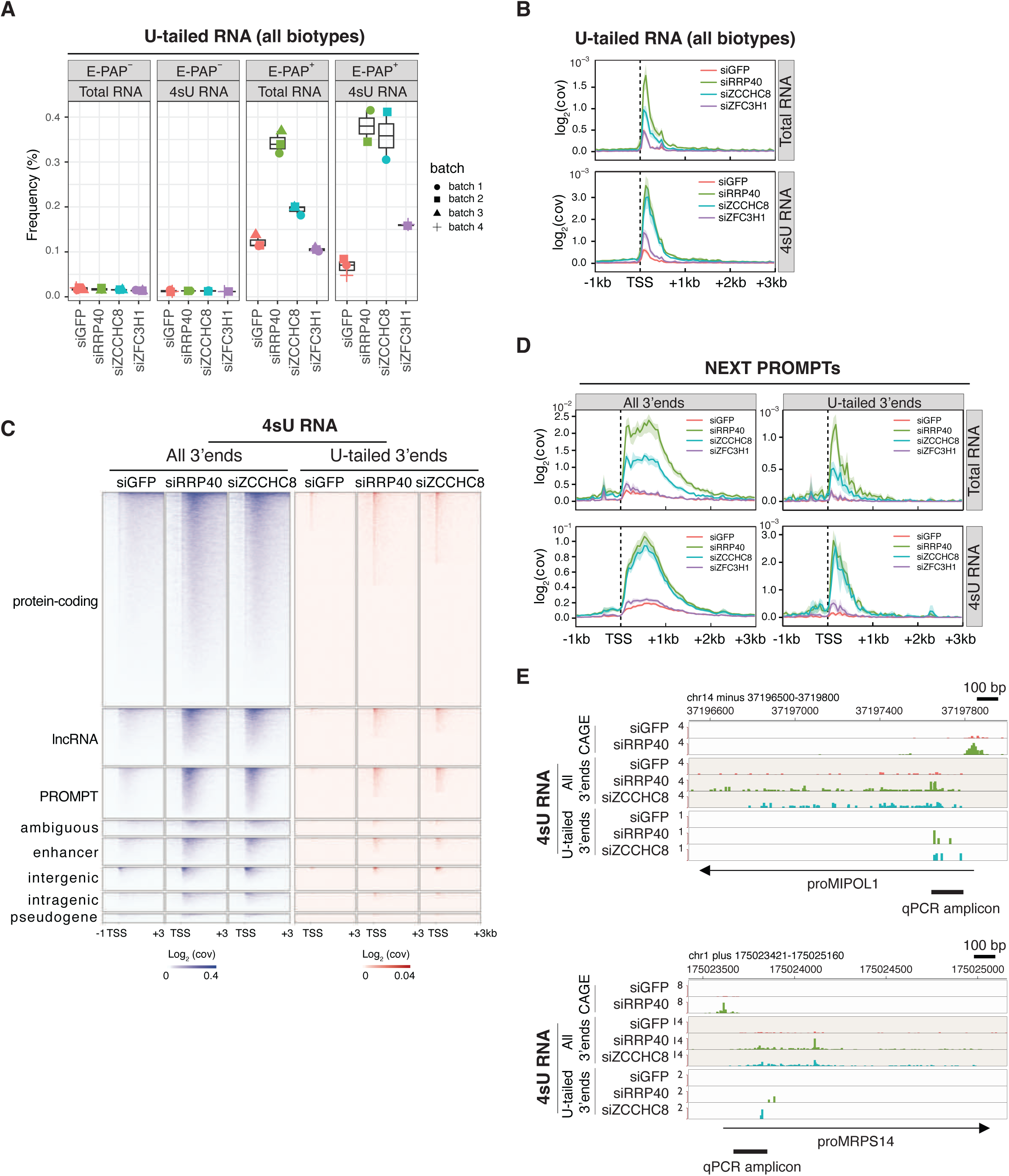
The human transcriptome pervasively generates short 3’end uridylated RNAs. **(A)** Frequencies of U-tailed RNA as defined by the ratio of reads with non-templated Us (3≤ U≤9) and total RNA reads. Data samples from cells transfected with siRNAs against GFP (siGFP, control), RRP40 (siRRP40), ZCCHC8 (siZCCHC8) or ZFC3H1 (siZFC3H1) are shown (GEO: GSE137612, ^36^). Data from samples without (E-PAP^−^) or with (E-PAP^+^) *in vitro* E-PAP treatment are shown separately. The displayed data constitute three biological replicates of total RNA, and two biological replicates of 4sU RNA from each of the depletion conditions, except for siGFP/4sU-RNA which was included in triplicate. Sequencing batch numbers are shown on the right. **(B)** Aggregate plots of reads of cellular U-tailed RNAs from total- and 4sU-RNA samples as indicated. The log_2_ data coverage (log_2_(cov)) was plotted from regions encompassing -1kb upstream to 3kb downstream of annotated transcription start sites (TSSs) ^50^. Mean values and 90% confidence intervals are shown. **(C)** Heatmaps of transcript end sites (TESs) of all RNAs (‘All 3’ends’) or U-tailed RNAs (‘U-tailed 3’ends’) from -1kb to 3kb regions relative to the TSSs of the indicated RNA biotypes as defined in ^50^. Data from 4sU RNA samples are shown. All heatmaps were sorted in descending order according to the log_2_(cov) values in each depletion sample. **(D)** Metagene profiles as in (B), but showing ’All 3’ends’ (left) or ‘U-tailed 3’ends’ (right) of NEXT-sensitive PROMPTs split by total (upper) and 4sU (lower) RNA samples. **(E)** Genome browser views of ‘All 3’end’ and ‘U-tailed 3’end’ data from the indicated 4sU RNA samples. Normalized and averaged signals are shown. PROMPT names (*i.e.* proMIPOL1 and proMRPS14) are shown at the bottom of each screenshot. RNA 5’end data indicating PROMPT TSSs were derived from cap analysis of gene expression (CAGE) data ^114^. qRT-PCR amplicons used for downstream analyses are indicated. See also Figure S1.

To analyze for a potential RNA biotype specificity underlying the observed U-tailing, we generated biotype-stratified heatmaps of the 3’end seq data. Distinctly, increased U-tail levels were observed for all RNA biotypes in exosome- and NEXT-depletion conditions, and in both total- and 4sU-RNA samples (Figure 1C and S1D). However, compared to the general upregulation of RNA (‘all 3’ends’), U-tailing was predominantly present on shorter RNAs. In this relation, we note that transcripts < 100nt were excluded from our libraries due to the utilized RNA purification method, which therefore are likely to underestimate RNA U-tail frequencies. Focusing on well-defined NEXT-sensitive PROMPTs ^36^, a similar pattern of upregulation of short RNAs with U-tails upon exosome- and NEXT-depletion was evident (Figure 1D). Like the general 3’end heterogeneity of NEXT-sensitive PROMPTs^36^, U-tailed transcripts also presented diverse 3’ends (Figure 1E and S1E).

Taking our analysis together, we conclude that RNA U-tailing is pervasive and generally more abundant upon disruption of the NEXT/exosome pathway. Hence, when NEXT activity is low, RNAs appear, which can both be uridylated and adenylated ^36^.

### Short excess NEXT substrates are uridylated by TUT4/7 and degraded by cytoplasmic exonucleases

In human cells, RNA terminal uridylyl transferase activity is mainly mediated by the TUT4 and TUT7 enzymes, which uridylate RNAs redundantly, but also with substrate preference, to convey RNA processing or degradation ^57,58,62,63^. With this in mind, we wondered whether TUT4 and/or TUT7 could uridylate NEXT-sensitive RNAs and control their abundance in the absence of functional nuclear exosome- or NEXT-complexes. Hence, siRNAs targeting TUT4 and TUT7 individually, or in combination, were transfected into HeLa cells, expressing RRP40 with a mini-auxin-inducible degron tag (RRP40-mAID), enabling rapid protein depletion upon the addition of indole-3-acetic acid (IAA/auxin) ^64^ (Figure S2A and S2B). TSS-proximal PCR amplicons for 8 NEXT-sensitive PROMPTs were designed, based on our RNA 3’end seq data (see examples in Figure 1E), and used to monitor the relative RNA abundance by qRT-PCR. Given the heterogeneity of the interrogated substrates, a mixture of random hexamer- and anchored dT_20_ (dT_20-_VN)-primers were utilized for cDNA synthesis to convert all RNA isoforms for subsequent qPCR analysis. Combined depletion of TUT4 and TUT7 (TUT4/7) mildly affected PROMPT levels, which was exacerbated when RRP40 was co-depleted by auxin addition (Figure S2C). A similar trend was observed in a background of ZCCHC8-mAID cells upon their induced ZCCHC8 depletion (Figure S2D, S2E, and S2F). We interpret these results to indicate that NEXT and the nuclear exosome normally clears most of the measured substrates, but upon NEXT/exosome inactivation TUT4/7 contribute to suppressing RNA levels.

To more precisely pinpoint how TUT4/7 depletions affected tailing of the interrogated RNAs, we devised a method to semi-selectively measure U-tailed RNA frequencies. Briefly, total cellular RNA was mixed with two *in vitro* transcribed spike-in RNA species: i) spike-in RNA 1^#^; a mixture of 6 RNA isoforms containing 0 to 5 Us at their 3’ends, and ii) spike-in RNA 2^#^; a negative control RNA devoid of 3’terminal Us (Figure S2G). This RNA cocktail was polyadenylated by E-PAP and subjected to reverse transcription using a dT_20-_A_4_ primer for detection of U-tailed RNA (>3U) by subsequent qPCR analysis. The dT_20-_A_4_ primer was tested together with other primers (see details in Methods) and chosen due to its high specificity for spike-in 1^#^ RNA (observed U-tailed RNA frequency of 22% and close to the theoretical value of 20%) and low detection of the spike-in 2^#^ control (∼2%) (Figure S2H). In parallel, levels of all RNA isoforms were measured from cDNA synthesized with the unselective dT_20-_VN primer, which allowed the calculation of U-tailed RNA frequencies. Gratifyingly, using this E-PAP-based approach to measure levels of all RNA isoforms (Figure 2A) recapitulated our results using random hexamer- and dT_20-_VN-primers (Figure S2C). Furthermore, a significant decrease of U-tailed RNA frequencies was detected upon TUT4/7 depletion, consistent with the uridylyltransferase activities of these enzymes (Figure 2B). As expected, similar phenotypes were observed in ZCCHC8-mAID cells upon depleting TUT4/7 (Figure 2C and 2D).

**Figure 2.**
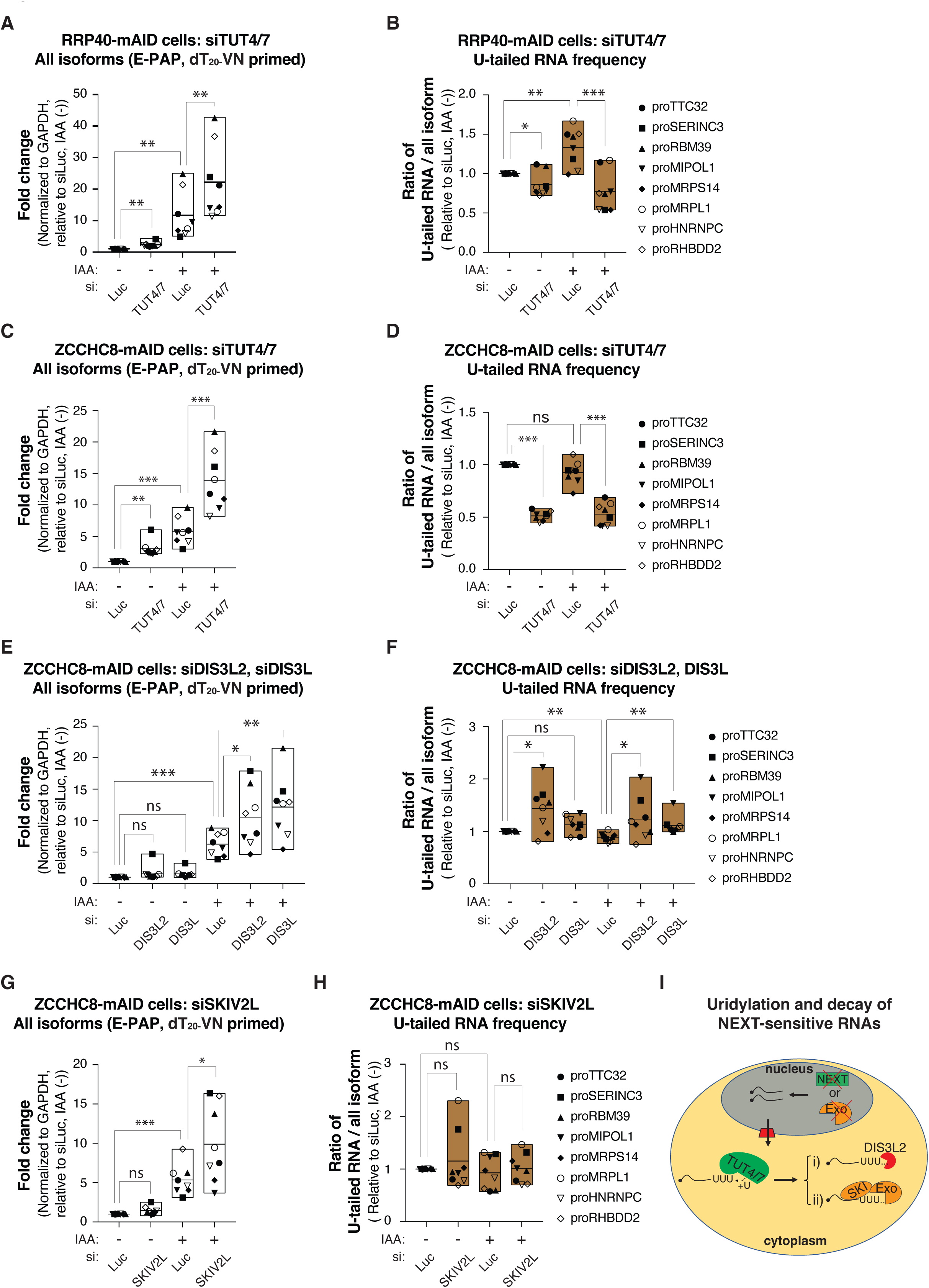
Cytoplasmic 3’-5’ exoribonucleases degrade NEXT-sensitive RNAs uridylated by TUT4/7. **(A)** Boxplots of qRT-PCR analysis of NEXT-sensitive RNAs from RRP40-mAID cells, transfected with siLuc or siTUT4/7, and treated with auxin (+), or not (-), as indicated. cDNA was synthesized using a dT_20-_VN primer after E-PAP treatment of total RNA. Individual levels of the 8 RNAs (see right part of Figure 2B) are represented by separate symbols with the mean value of three biological replicates shown. The line inside each box represents the mean value of the 8 targets. qRT-PCR data were normalized to GAPDH (reference RNA) and plotted relative to the control sample (siLuc, IAA (-)). Statistical analysis was performed using two-sided t tests. * p< 0.05; ** p < 0.01; *** p< 0.001; ns, not significant. Similar statistics were employed for subsequent analyses. Note that boxplots, which display measurements of all RNA isoform levels (‘All isoforms’) after E-PAP treatment and dT_20_-NV priming condition, are consistently represented in white color. **(B)** Boxplots of qRT-PCR analysis of U-tailed RNA frequencies as described in (Figure S2G). Note that boxplots, which display measurements of U-tailed RNA frequencies, are consistently represented in brown color. **(C)** Boxplots of qRT-PCR analysis as in (A), but employing ZCCHC8-mAID cells. **(D)** Boxplots of qRT-PCR analysis of U-tailed RNA frequencies as in (B), but employing ZCCHC8-mAID cells. **(E)** Boxplots of qRT-PCR analysis as in (C), but employing siDIS3L2 or siDIS3L siRNAs as indicated. **(F)** Boxplots of U-tailed RNA frequencies analysis as in (D), but employing siDIS3L2 or siDIS3L siRNAs as indicated. **(G)** Boxplots of qRT-PCR analysis as in (C), but employing siSKIV2L siRNA as indicated. **(H)** Boxplots of U-tailed RNA frequencies analysis as in (D), but employing siSKIV2L siRNA as indicated. **(I)** Diagram summarizing the proposed uridylation and decay pathway. See also Figure S2.

How do these U-tailed RNAs get degraded? As TUT4/7 localize to the cytoplasm of human cells ^54,65^, we focused on two possible cytoplasmic decay pathways; i) 3’-5’ RNA decay by DIS3L2, and ii) 3’-5’ decay by the cytoplasmic exosome, represented by DIS3L. As RRP40 is the subunit of both nuclear and cytoplasmic exosomes, individual siRNA-mediated depletion of these RNA exonucleases was carried out in ZCCHC8-mAID cells (Figure S2I). Scrutinizing levels of all RNA isoforms established that depletion of either DIS3L2 or DIS3L increased transcript levels in ZCCHC8 depletion conditions (Figure 2E). Moreover, and in contrast to TUT4/7 depletion conditions, the frequencies of U-tailed RNAs were slightly increased (Figure 2F). To substantiate a role of the cytoplasmic exosome, we also depleted SKIV2L (Figure S2J), a component of its accessory SKI complex ^9,66^. Consistent with the effect of DIS3L depletion, SKIV2L depletion also increased transcript levels (Figure 2G), without changing U-tailed RNA frequencies as U-tailed RNA levels increased correspondingly (Figure 2H).

Taking our results together, we conclude that NEXT-sensitive PROMPTs, upon escaping NEXT-mediated nuclear exosome decay, can be exported to the cytoplasm and uridylated by TUT4/7 (Figure 2I). Uridylated RNAs are then degraded by DIS3L2 or by the cytoplasmic exosome, providing a back-up system for when nuclear RNA decay is compromised. While an involvement of DIS3L2 in degrading U-tailed RNA is unsurprising ^15,63,67,68^, the cytoplasmic exosome has not previously been implicated in U-tailed RNA degradation. However, it has been reported to degrade U-tailed nonsense-mediated mRNA decay (NMD) intermediates ^69^ and SKIV2L can bind U-tailed RNA ^66^. Thus, decay of U-tailed RNA by the cytoplasmic exosome could be a more general mechanism.

### PHAX exports short excess NEXT substrates for cytoplasmic uridylation

We next addressed how short excess NEXT substrates end up in the cytoplasm after escaping nuclear decay. Since the PHAX protein is involved in the export of short capped RNA, such as UsnRNA, short ncRNA and the H2AX RNA, due to its affinity for RNA < 300 nt ^20–23^, we interrogated this possibility by depleting cellular PHAX using siRNA. As observed for siTUT4/7, co-depletion of PHAX with RRP40 significantly increased NEXT-sensitive PROMPT levels (Figure 3A and S3A), while markedly decreasing U-tailed RNA frequencies (Figure 3B). Similar tendencies were observed upon co-depletion of PHAX with ZCCHC8 (Figure 3C, 3D and S2I). We note that here PHAX depletion on its own impacted RNA levels, which likely attributes to lowered ZCCHC8 levels upon mAID-tagging ^64^. Nevertheless, these data, thus, supported the idea that PHAX is involved in the nuclear export of short excess NEXT substrates to make them accessible for uridylation in the cytoplasm.

**Figure 3.**
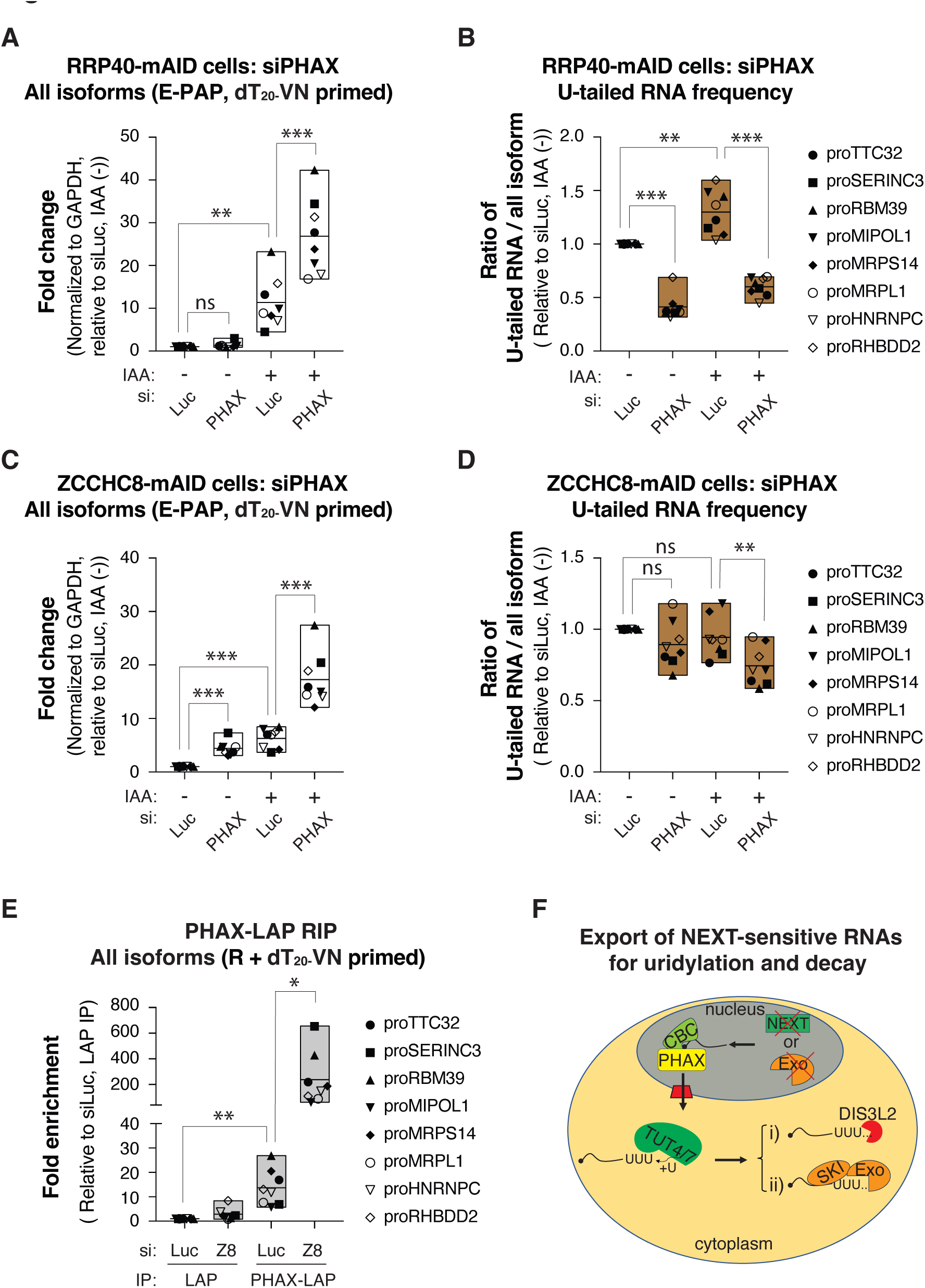
PHAX exports short NEXT-sensitive RNAs. **(A)** Boxplots of qRT-PCR analysis as in Figure 2A, but depleting PHAX. **(B)** Boxplots of U-tailed RNA frequencies as in Figure 2B, but depleting PHAX. **(C)** Boxplots of qRT-PCR analysis as in (A), but employing ZCCHC8-mAID cells. **(D)** Boxplots of U-tailed RNA frequencies as in (B), but employing ZCCHC8-mAID cells. **(E)** Boxplots of qRT-PCR analysis of the indicated RNA levels from IP samples from Figure S3B. The values of the control sample (siLuc, LAP IP) were set to 1. cDNA for qPCR analysis was synthesized by a mix of random hexamer- and dT_20-_VN-primers (R+dT_20-_VN). Data show the mean values of two biological replicates. Note ‘Z8’ is short for ZCCHC8. **(F)** Diagram summarizing the proposed RNA export, uridylation and decay pathways. See also Figure S3.

To evaluate PHAX binding to the investigated transcripts, we conducted RNA immunoprecipitation (RIP) using a GFP antibody to precipitate PHAX from HeLa cells, stably expressing a ‘localization and affinity purification’ (LAP)-tagged version of PHAX (PHAX-LAP) ^70,71^. As expected, given the recruitment of PHAX to RNA via the CBC ^21^, a specific interaction between PHAX-LAP and the CBP20 and CBP80 proteins could be confirmed, which was not affected by ZCCHC8-depletion (Figure S3B). More importantly, the PHAX-LAP IP enriched for NEXT substrates in the control (siLuc) condition, which was further increased upon ZCCHC8 depletion (Figure 3E), presumably due to increased substrate availability (Figure S3C). We suggest that in the absence of NEXT, its short RNA substrates bind to PHAX, which mediates their nuclear export, providing for cytoplasmic uridylation-coupled degradation (Figure 3F).

### A fraction of longer excess NEXT substrates is polyadenylated and exported to the cytoplasm

Inspired by short excess NEXT substrates, utilizing nuclear export and cytoplasmic U-tailing for their decay, we wondered whether A-tailed excess NEXT substrates would employ a similar backup mechanism, in addition to their demonstrated handover for PAXT-mediated decay in the nucleus ^36^. Our RNA 3’end seq data revealed that A-tailed substrates, accumulating upon ZCCHC8 depletion, were generally longer than their U-tailed counterparts (Figure 4A, compare to Figure 1D, right panel). Moreover, and as previously shown ^36^, levels of A-tailed substrates were increased upon co-depletion of ZCCHC8 with the PAXT component ZFC3H1 (Figure 4A).

**Figure 4.**
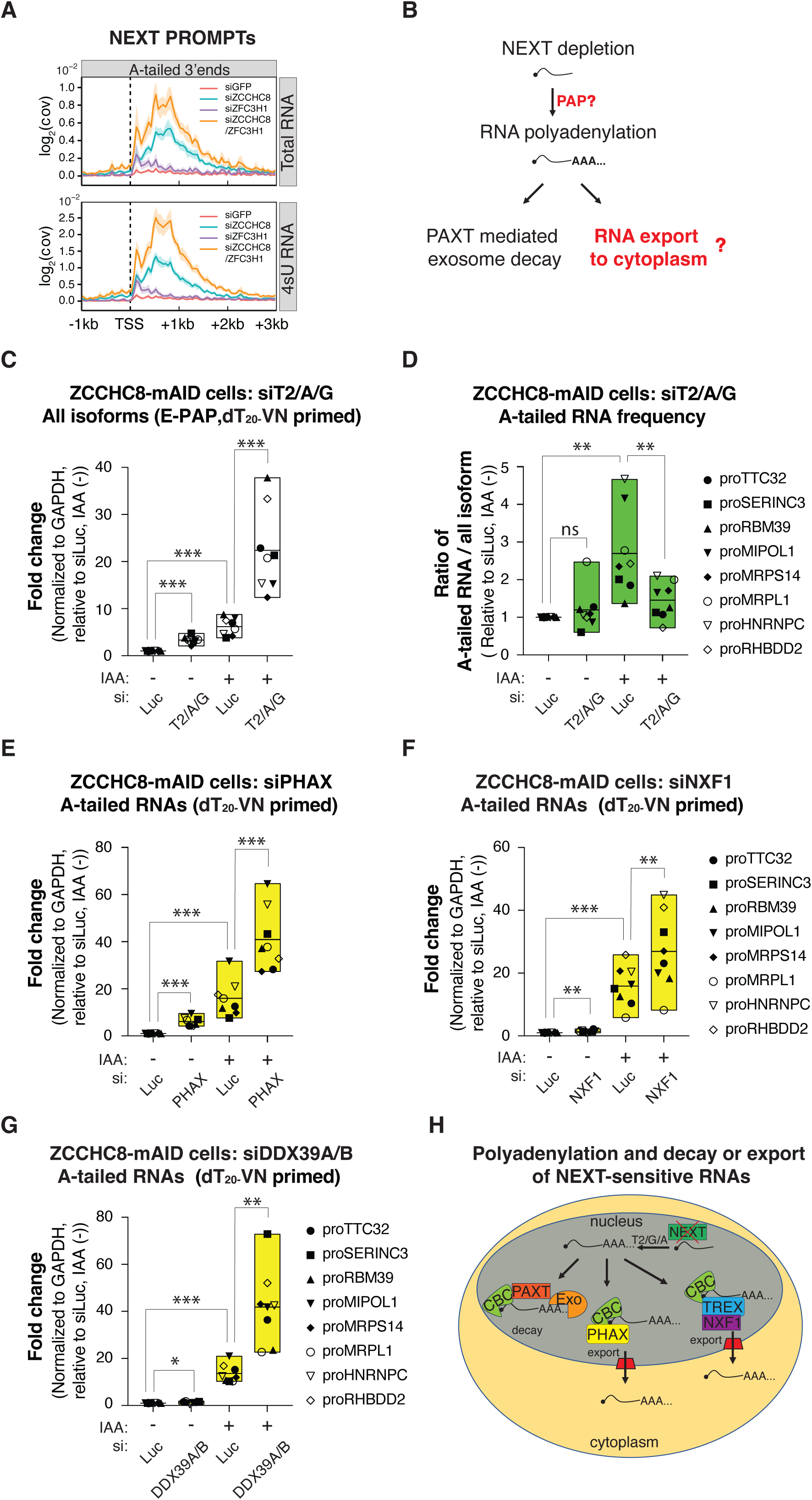
NEXT-sensitive RNAs can be polyadenylated and exported to the cytoplasm. **(A)** Metagene profiles as in Figure 1D, but showing ‘A-tailed 3’ends’ of NEXT-sensitive PROMPTs from the indicated depletion samples, displaying total- (upper) or 4sU- (lower) RNA. Data from no E-PAP treatment samples were used (data in GEO: GSE137612, ^36^). **(B)** Schematic diagram showing the proposed nuclear polyadenylation of excess NEXT substrates followed by their nuclear decay or export. Red text and question marks indicate unresolved processes. **(C)** Boxplots of qRT-PCR analysis as in Figure 3C, but depleting TENT2/PAPOLA/PAPOLG (T2/A/G). **(D)** Boxplots of qRT-PCR analysis of samples from (C) but measuring A-tailed RNA frequencies as depicted in Figure S4F. Note that this boxplot, which displays measurements of A-tailed RNA frequencies, is represented in green color. **(E)** Boxplots of qRT-PCR analysis of A-tailing levels of RNAs from Figure 3C. To obtain A-tailed RNA, total RNA samples were reverse transcribed with a dT_20-_VN primer before qPCR analysis. Note that boxplots, which display measurement of A-tailed RNA levels (‘A-tailed RNAs’), are consistently represented in yellow color. **(F)** Boxplots of qRT-PCR analysis as in (E), but depleting NXF1. **(G)** Boxplots of qRT-PCR analysis as in (E), but depleting DDX39A/B. **(H)** Diagram summarizing the proposed PAXT-mediated exosome decay or nuclear export of A-tailed excess NEXT substrates. See also Figure S4.

Despite the fact that pA^-^ NEXT-substrates can be polyadenylated, the responsible poly(A) polymerase(s) (PAPs) remained unclear as did the probability of whether a fraction of these A-tailed RNAs could be subjected to nuclear export (Figure 4B). To address the first question, we depleted canonical PAPs, PAPOLA and PAPOLG, as well as their non-canonical, but nuclear, counterparts TENT1, TENT4A/TENT4B (TENT4A/B) and TENT2 (Figure S4A, S4B and S4C). Individual depletion of TENT2, PAPOLA and PAPOLG mildly increased NEXT substrate levels in the ZCCHC8 depletion condition, but no effect was observed upon TENT1 or TENT4A/B depletion (Figure S4D). To address any possible redundancy, TENT2, PAPOLA and PAPOLG were co-depleted (Figure S4C and S4E), which led to more robust RNA accumulation (Figure 4C). This was mirrored by a decreased RNA A-tailing frequency (Figure 4D and S4F). We therefore conclude that A-tailing of excess NEXT substrates can be conducted redundantly by TENT2, PAPOLA and PAPOLG.

To next explore the possibility of nuclear export of A-tailed RNAs, their levels were examined by co-depleting candidate export factors with ZCCHC8. PHAX co-depletion increased A-tailed RNA levels (Figure 4E), which was consistent with its binding to these substrates as demonstrated by RIP experiments (Figure S4G and S4H). We noted, however, that PHAX binding here was less prominent when compared to its interaction with all RNA isoforms (compare Figure S4H and 3E, note Y-axis scales). Given that most A-tailed RNAs are believed to be exported by NXF1 with the assistance of the TREX complex ^18,19,72^, we co-depleted NXF1 or the TREX component DDX39A/B (DDX39A/DDX39B) along with ZCCHC8 (Figure S4I and S4J). Both depletions yielded increased levels of A-tailed RNA (Figure 4F and 4G), suggesting that these excess NEXT substrates can also be exported by the TREX-NXF1 pathway, possibly because the nuclear PAXT/exosome decay pathway becomes saturated.

Taking the data together, we suggest that excess NEXT substrates can be redundantly polyadenylated by TENT2, PAPOLA and PAPOLG, leading to PAXT-mediated nuclear decay or PHAX- or TREX-NXF1 mediated export (Figure 4H).

### Exported A-tailed excess NEXT substrates are degraded in a translation-dependent manner

The observed increase in A-tailed excess NEXT substrates, upon export factor depletion, suggested their removal in the cytoplasm. To interrogate by which mechanism(s) such decay might be carried out, we examined A-tailed substrate levels from ZCCHC8-mAID samples co-depleted for the exosome exoribonuclease DIS3L, or DIS3L2 (Figure S2I). While DIS3L2 depletion, in line with its uracil-specificity ^15,67^, did not affect A-tailed RNA levels (Figure S5A), these increased upon DIS3L depletion (Figure 5A). Consistent with an involvement of the cytoplasmic exosome, depleting SKIV2L (Figure S2J) also increased levels of excess A-tailed NEXT substrates (Figure 5B). Moreover, disrupting both nuclear and cytoplasmic decay by co-depleting ZFC3H1 and SKIV2L (Figure S2J) further increased A-tailed RNA levels (Figure 5B), suggesting that the two decay pathways cooperate to clear these substrates.

**Figure 5.**
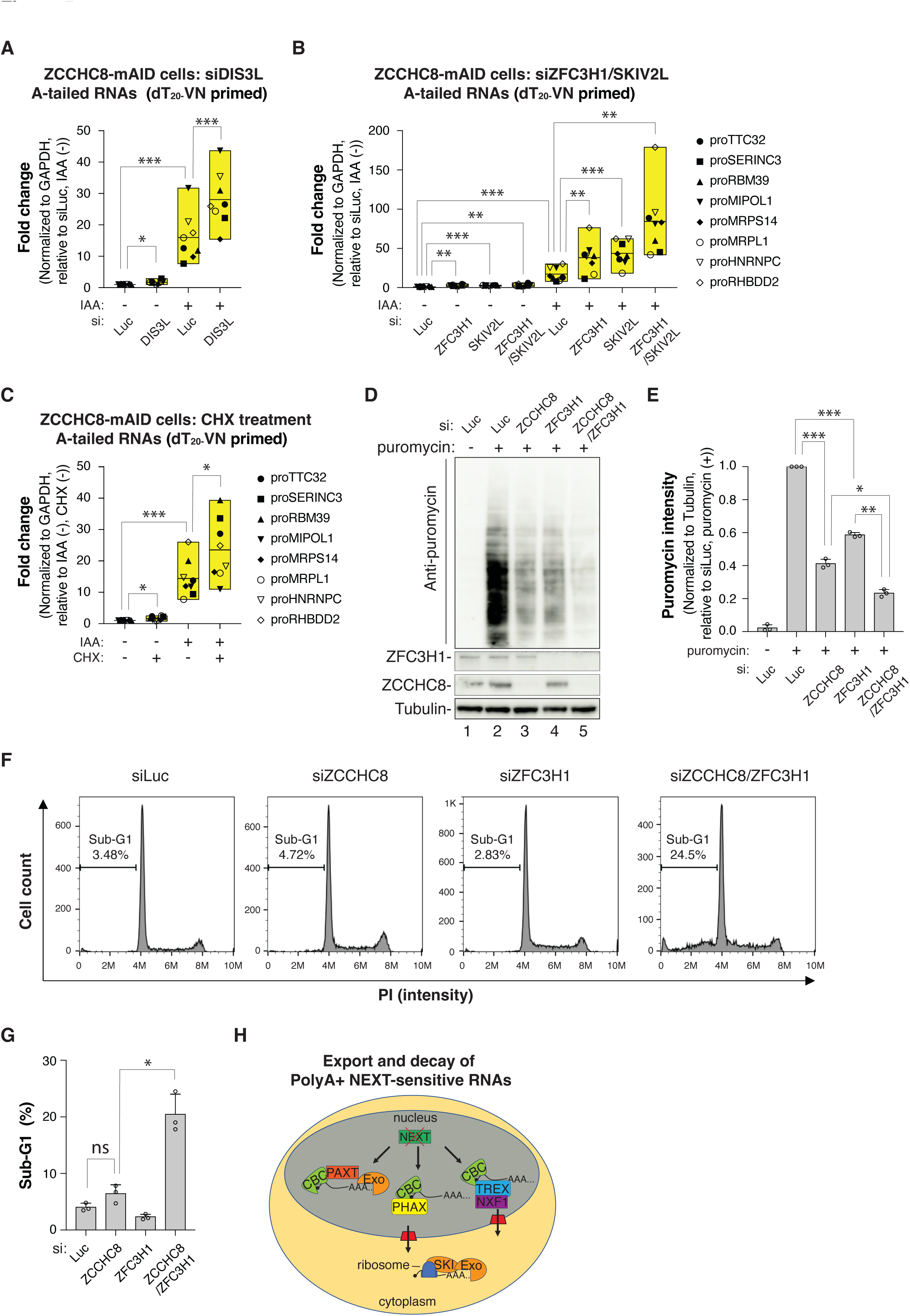
SKI/exosome degraded the A-tailed NEXT-sensitive RNAs in cytoplasm. **(A)** Boxplots of qRT-PCR analysis as in Figure 4E, but employing the indicated samples from Figure S2I. **(B)** Boxplots of qRT-PCR analysis as in (A), but employing the indicated samples from Figure S2J. **(C)** Boxplots of qRT-PCR analysis as in (A), but treating cells without (-) or with (+) 10 ug/ml cycloheximide (CHX) for 3 hours. **(D)** Western blotting analysis assessing puromycin incorporation in cells upon the indicated siRNA-mediated protein depletions. HeLa cells were siRNA transfected for 72 h, and treated with 5 ug/ml puromycin for 30 min before harvesting for western blotting analysis. Western membranes were probed with antibodies towards puromycin, ZCCHC8, ZFC3H1 and Tubulin as a loading control. A representative experiment from biological triplicate data is shown. **(E)** Quantification of triplicate experiments from (D). Puromycin signal intensities were normalized to those of Tubulin and with the control sample (siLuc, puromycin (+) set to 1. Biological triplicate data are shown with each circle representing single replicate values. **(F)** Flow cytometry analysis of the cell cycle distribution of cells subjected to the indicated siRNA-mediated protein depletion. The percentages of sub-G1 cells are indicated. A representative experiment from biological triplicate data is shown. **(G)** Quantification sub-G1 population data from (F). Biological triplicate data are shown with each circle representing single replicate values. **(H)** Diagram summarizing the proposed nuclear PAXT/exosome decay, export and cytoplasmic SKI/exosome decay of A-tailed excess NEXT substrates. See also Figure S5.

To investigate whether cytoplasmic SKI/exosome degrades excess A-tailed NEXT substrates in a translation-dependent or -independent manner, we treated ZCCHC8-mAID cells with the translation elongation inhibitor cycloheximide (CHX). This led to increased levels of A-tailed RNA (Figure 5C), suggesting translation-dependent turnover, as also observed for other cytoplasmic exosome substrates ^9,66,73^. Interestingly, however, although DIS3L dually participates in A-tailed and U-tailed RNA decay (Figure 2F and 5A), turnover of the latter was not CHX-sensitive (Figure S5B).

The engagement of translation in the turnover of cytoplasmic A-tailed excess NEXT substrates might occupy available ribosomes and, in turn, affect normal translational output. To examine this possibility, we estimated protein synthesis by labeling nascent peptides with puromycin, which causes premature chain termination by its covalent incorporation into the growing peptide chain measurable by anti-puromycin western blotting analysis ^74–76^. As our ZCCHC8-mAID cells harbor the puromycin resistance gene, we employed regular HeLa cells and instead depleted ZCCHC8 and/or ZFC3H1 by siRNA transfection (Figure 5D, bottom panels). Individual depletion of these factors led to decreased puromycin levels, which was further lowered by ZCCHC8/ZFC3H1 co-depletion (Figure 5D, top panel and 5E). Thus, excess RNA released from the nucleus appears to affect global protein synthesis, possibly by occupying the translation machinery. Consistent with such importance of ZCCHC8 and ZFC3H1 in maintaining normal protein synthesis, a lower cell confluency was observed in this co-depletion sample (data not shown). This indicated increased cell death, which was validated by flow cytometry analysis, revealing an increased sub-G1 population accompanied by a decreased G0/G1 population (Figure 5F, 5G and S5C).

Taking these results together, we suggest that A-tailed excess NEXT substrates can either be degraded in the nucleus by the PAXT/exosome pathway or in the cytoplasm by the SKI/cytoplasmic exosome complex in a translation-dependent manner (Figure 5H). Collectively, this possibly protects against an overpowering of physiological translation and promotes cell survival.

### Fail-safe decay pathways target NEXT substrates upon DNA damage

Our results so far revealed fail-safe removal of NEXT substrates under conditions where this nuclear exosome adaptor was artificially depleted. To address whether the identified NEXT back-up systems would also respond to other conditions affecting NEXT activity, like DNA damage where RBM7 is phosphorylated and in turn lowers its affinity towards decay substrates ^41–43^, we challenged cells with the DNA damage-inducing agent 4-nitroquinoline 1-oxide (4-NQO). As expected, 4-NQO administration increased levels of the previously interrogated ‘All isoforms’ (Figure 6A, compare siLuc 4-NQO (-) and (+)), consistent with NEXT inactivation ^41,43^. Moreover, depleting SKIV2L or DIS3L2 in 4-NQO treated cells (Figure S6) further increased RNA levels (Figure 6A), while leaving U-tailed RNA frequencies unchanged or slightly increased (Figure 6B). Finally, depleting ZFC3H1 or SKIV2L, or both, increased A-tailed RNA levels in 4-NQO treated cells (Figure 6C) and combining siZFC3H1 with 4-NQO treatment induced a significantly increased of sub-G1 populations (Figure 6D and 6E). We therefore conclude that fail-safe decay is also operational upon DNA damage.

**Figure 6.**
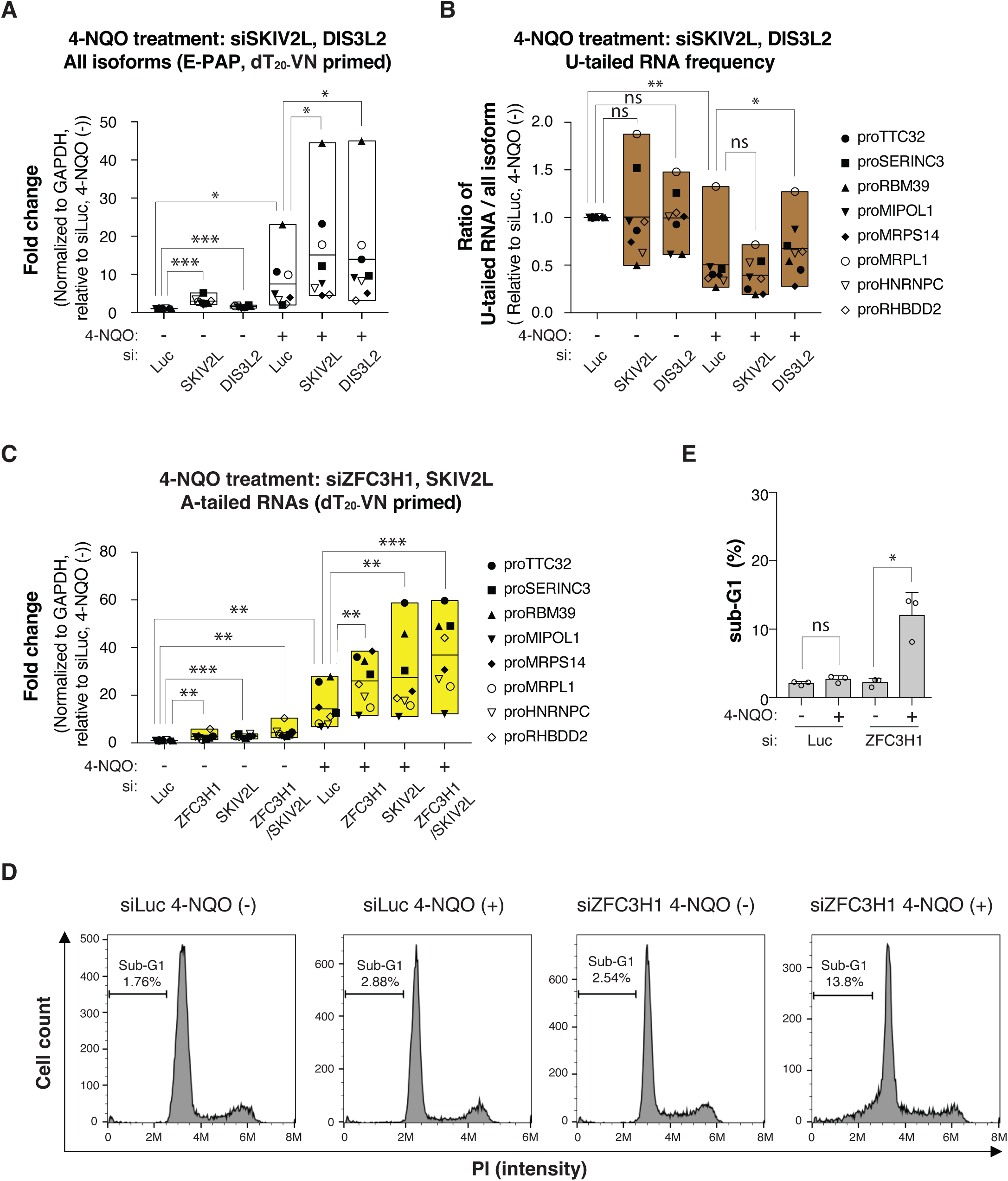
Fail-safe pathways degrade NEXT substrates upon DNA damage. **(A)** Boxplots of qRT-PCR analysis as in Figure 2A but employing the indicated depletion samples treated with 5 µM 4-NQO (+), or not (-) for 4 hours. **(B)** Boxplots of qRT-PCR analysis as in (A), but measuring U-tailed RNA frequencies. **(C)** Boxplots of qRT-PCR analysis as in Figure 3A but employing the indicated depletion samples with or without 4-NQO treatment. **(D)** Flow cytometry analysis as in Figure 5F, but employing the indicated samples with or without 4-NQO treatment. **(E)** Quantification of sub-G1 cell populations from (E). Biological triplicate data are shown with each circle representing single replicate values. See also Figure S6.

## Discussion

Decay systems rid cells of aberrant and excess transcripts to maintain RNA homeostasis. This encompasses major RNA decay pathways, across the nucleus and the cytoplasm ^3,4^, but how these interrelate to compensate for one another has not been thoroughly explored. In one example from higher eukaryotes, we have previously described that the inefficient removal of pA^-^ RNA substrates by the nuclear exosome, due to depletion of its NEXT adaptor complex, may trigger their post-transcriptional 3’end adenylation and subsequent handover to PAXT-mediated nuclear decay by the exosome ^36^. Earlier data from budding yeast have demonstrated that handover of excess RNA can also occur across cellular compartments as transcripts escaping nuclear quality control are exported and abundantly targeted by the cytoplasmic nonsense mediated decay (NMD) pathway ^77^. In the present study, we now expand this principle by describing NEXT pathway backup mechanisms, relying on RNA 3’end tailing and nuclear export, albeit in sequentially opposite orders, to reach their consequential cytoplasmic decay destinies (Figure 3F and 5H). The need for cells to be able to efficiently handle excess NEXT substrates is apparent given the wide variety of such substrates, including PROMPTs ^30^, eRNAs ^78^, 3’extended snRNAs, snoRNAs, histone RNAs and telomerase RNAs ^64,78–80^ as well as prematurely terminated RNAPII products from within protein-coding genes ^47,50–52^ and from within transposable elements ^81,82^. We therefore suggest that primary decay by NEXT/exosome and compensation by backup systems cooperate to restrict these non-productive RNAs, enabling cells to handle extrinsic challenges, such as the DNA damage condition imposed in this study, ultimately promoting cell survival.

The discovered backup mechanisms involve RNA 3’end tailing, implying that at least some RNAs with naked 3’ends are not optimal decay substrates after their escape from initial NEXT targeting. In previously reported cases, NEXT complex activity occurs in tight conjunction to the prior processes of transcription termination ^50,52,82^, pre-mRNA splicing ^64,83^ or microprocessor cleavage ^84^ and in select examples a direct physical interaction between RNA substrate production- and decay-systems have been demonstrated ^51,52,84^. Thus, while conventional turnover of NEXT substrates is typically a coupled processes that accepts an unmodified RNA 3’end, their ‘pick up’ by alternative decay pathways, described here, rely on prior 3’end tailing.

In the instance of the shorter fraction of our interrogated excess NEXT substrates, cytoplasmic 3’end uridylation proceeds decay by DIS3L2 or the SKI-associated exosome. RNA U-tailing is generally considered a mark for complete RNA decay, although transcript processing might also ensue, *e.g.* uridylation of U6 snRNA ^85–87^ and mono-uridylation of *Let-7* miRNA ^88–90^. Despite the fact that non-templated U-tails are found on a range of RNA substrates, and that the operational uridylation enzymes TUT4 and TUT7 have been identified, the mechanisms underlying TUT4/7 recruitment to RNA 3’ends have only been delineated in a few individual cases. In one example, the *Let-7* miRNA binding protein Lin28 directly interacts with TUT4/7 in embryonic stem cells ^91,92^ and in another, TUT4/7 targeting to transcripts, deriving from LINE-1 retrotransposable elements, is reported to occur via the MOV10 protein ^54^. Since DIS3L2 was found to bind structured RNAs, RNA secondary structure has been suggested to directly, or indirectly through RNA binding proteins, recruit TUT4/7 ^17^. However, the diverse set of DIS3L2 and SKI-exosome substrates reported in the present study suggest that a more general feature of these transcripts might be recognized. Given the shortness of the identified U-tailed excess NEXT substrates, RNA size might be relevant for TUT4/7 recruitment. Perhaps the difficulty of these RNAs to associate with ribosomes enhances the likelihood of TUT4/7 targeting. This could direct SKIV2L to the U-tailed RNA, either directly ^66^, or via an intermediary protein, as seen for fungus-specific SKA1 protein ^93^. In an alternative, but not mutually exclusive, scenario the nuclear history of these transcripts might impact the involved cytoplasmic processes. In the absence of active NEXT-targeting, the competing process of PHAX-mediated nuclear export prevails ^37,70^, possibly enabling the resident RNA to be subjected to cytoplasmic 3’end modification as is the case for the conventional UsnRNA PHAX-export substrates ^94,95^. However, different from the productive 3’end processing and RNP assembly of snRNA prior to their nuclear re-import, excess NEXT substrates are tailed and subjected to removal. Which mechanism(s) might discriminate these productive and destructive pathways remains an interesting matter for further investigation.

While RNA U-tailing is primarily involved in destructive pathways, A-tailing is widely employed for both productive and destructive purposes. Conventional 3’end polyadenylation by the cleavage and polyadenylation (CPA) complex largely occurs co-transcriptionally and is critical for further RNA processing, stability, nuclear export and translation ^96–99^. In a countering destructive pathway, the RNA polyA tail and its nuclear pA-binding protein (PABPN1) can direct transcripts for decay by the PAXT/exosome pathway ^30–32,36,39,40,100–103^. Additionally, short polyA tails, added by the mammalian Trf4/Air2/Mtr4 polyadenylation (TRAMP) complex promote the turnover of rRNA processing byproducts through the nuclear exosome ^30,104^. In an apparent blend of these pathways, excess NEXT substrates can be A-tailed in the nucleus in preparation for their PAXT-mediated destruction (^36^ and the present study). We show that such A-tailing is redundantly conducted by the TENT2, PAPOLA and PAPOLG enzymes. Only scattered evidence exists on how this might occur. TENT2 is localized in both the nucleus and the cytoplasm ^105,106^, and was reported to adenylate cytoplasmic mRNA through recruitment by the cytoplasmic polyadenylation element-binding protein 1 (CEBP1) or the QKI7 KH domain-containing RNA binding (QKI-7) protein ^107–109^. Instead, both PAPOLA and PAPOLG are strictly nuclear and reportedly adenylate RNA in a CPA-dependent process coupled to pA site cleavage ^110,111^. Since NEXT substrates are originally produced as pA^-^ RNAs in a CPA-independent manner ^36^, their adenylation by TENT2, PAPOLA and PAPOLG is most likely to be posttranscriptional. However, as for 3’end uridylation, additional factors involved in attracting these pA polymerases to their pA^-^ RNA 3’ends remain to be defined.

In addition to their handover for PAXT-mediated decay, adenylated excess NEXT substrates can also be subjected to nuclear export. In fact, we show that such export is mediated by regular RNA export factors/complexes PHAX, TREX and NXF1, demonstrating that these unconventionally adenylated RNAs can still enter conventional export systems. This occurs in apparent competition with PAXT-mediated decay, shown here indirectly by the increased overwhelming of general translation upon co-depletion of the PAXT component ZFC3H1 (Figure 5D), and previously demonstrated directly by nuclear-cytoplasmic fractionation experiments ^38,103,112^. The major effects of ZCCHC8 and ZFC3H1 depletions on translation (Figure 5D) seemingly underscores the scale of cryptic RNAs normally removed in the nucleus and the central role of such decay in the maintenance of both RNA and protein homeostasis. It also suggests that most of these transcripts, upon entering the cytoplasm, does not produce appreciably amounts of protein but rather block ribosomes, preventing translation of proper mRNA. Cytoplasmic removal of the latter is mainly controlled by the translation-dependent and XRN1 mediated 5’-3’ RNA decay pathway ^66,113^. In contrast, the A-tailed excess NEXT substrates reported in this study are predominantly degraded in a translation-dependent process, depending on the SKI/exosome (Figure 5A and 5B), but not by XRN1 (data not shown). Since the SKI/exosome pathway primarily degrades specific RNAs as part of cytoplasmic quality control ^9,66^, particular features of adenylated excess NEXT substrates are likely to elicit their turnover. We suggest that these pertain to RNA characteristics, which are not evolved for efficient translation, including the short length of these transcripts, their RNP composition and/or the fact that their pA tails are not produced by the conventional CPA machinery.

Regardless the exact mechanisms, RNA 3’end tailing appears central for marking the RNA overflow, deriving from dampened NEXT-activity, for alternative downstream decay. This is eventually executed by pathways that are also operational in conventional RNA productive and destructive mechanisms and which underscores two main principles of RNA quality control: i) It is not carried out by specific systems, but rely on redundant conventional activities, and ii) these provide a network of specialized, but also cooperative pathways, enabling cells to maintain RNA homeostasis and to survive during stress.

## Acknowledgments

We thank Claudia Scheffler for expert technical assistance and members of the T.H.J. group for fruitful discussions. Søren Lykke-Andersen, Willian Garland and Andrii Bugai are thanked for their critical comments on the manuscript. The work was supported by the Novo Nordisk Foundation (ExoAdapt grant 31199) and the Danish Cancer Society (Grant no. R302-A17282). G.W. was supported by a Lundbeck Foundation Postdoc grant (R347-2020-2224).

## Author Contributions

G.W., and T.H.J. conceived the project. T.H.J supervised the project. G.W. performed all wet laboratory experiments. J.O.R. and M.S. performed computational analyses. G.W. and T.H.J wrote the paper. All authors read and approved the manuscript.

## Lead Contact

Further information and requests for resources and reagents should be directed to and will be fulfilled by the lead contact, Torben Heick Jensen.

## Methods

### Cell cultures and their manipulations

Wildtype, RRP40-mAID and ZCCHC8-mAID HeLa cells ^64^ were cultured in Dulbecco’s modified Eagle’s medium (DMEM) supplemented with 10% fetal bovine serum (FBS) and 1% penicillin/streptomycin (P/S). siRNA transfections were carried out using Lipofectamine RNAiMAX (Invitrogen) according to the manufacturer’s protocol. Cells were subjected to 20 nM siRNA treatment for 3 days in all depletion experiments, except for the siRNAs targeting NXF1 and DDX39A/B, which were restricted to 2 days due to severe cell death with prolonged depletion. For AID-mediated protein depletion, 750 µM IAA sodium salt (Sigma-Aldrich) was supplemented to cell culture medium for 6 hr before cell harvest. All siRNA sequences are listed in Table S1. For DNA damage induction, 5 µM of 4-NQO (Sigma-Aldrich) was added to cell culture media for 4 hr before cell harvest.

### Western blotting analysis

After harvesting, cells were counted using an automated cell counter (Invitrogen) and lysed with LDS Sample Buffer (Invitrogen) supplemented with Sample Reducing Agent (Invitrogen). Equal amounts of protein were loaded onto PAGE gels after sample denaturation at 95D for 15 min. Proteins were transferred to PVDF membranes, which were blocked with 5% skimmed milk in phosphate-buffered saline with 0.05% Tween 20 (PBS-T) for 1 hr at room temperature (RT), incubated with primary antibody diluted in PBS-T at 4℃ overnight and followed by washing 3×10 min with PBS-T. Membranes were then incubated with HRP-conjugated secondary antibody diluted in PBS-T for 1 hr at RT, followed by washing 3×10 min with PBS-T. SuperSignal West Femto HRP substrate (ThermoFisher Scientific) was applied to membranes and signals were detected with an ImageQuant 800 (Amersham). Utilized primary antibodies are listed in Table S2.

### RNA isolation and qRT-PCR analysis

RNA was extracted using TRIzol (Invitrogen) and treated with TURBO DNase (Invitrogen) according to the manufacturer’s protocol. For measuring ‘all isoform’ RNA levels, reverse transcription (RT) was carried out with SuperScript III reverse transcriptase (Invitrogen) using 1 µg RNA and a mixture of 20 pmol random hexamer- and 4 pmol dT_20-_VN-primers in a 20 µl reaction at 50 D according to the manufacturer’s protocol. Subsequent, qPCR was performed using Platinum SYBR Green qPCR SuperMix-UDG (Invitrogen) in a ViiA 7 Real-Time PCR machine (Life Technologies). When measuring ‘A-tailed’ RNA levels, qRT-PCR was carried out similarly, except that cDNA was synthesized using 20 pmol dT_20-_VN primers. Utilized qPCR primers are listed in Table S3.

### *In vitro* transcription of spike-in RNA

DNA templates for *in vitro* transcription were generated by PCR using a forward primer harboring a T7 promoter sequence and a reverse primer harboring 0-5 As at it 3’end. pUC19-ERCC-00136 (for spike in 1^#^) and ERCC-00136 (for spike in 2^#^) plasmids were used as PCR templates ^115^. *In vitro* transcription reactions were carried out using MEGAscript RNAi Kit (ThermoFisher Scientific) according to the manufacturer’s protocol. Subsequently, RNA was purified by phenol extraction and ethanol precipitation, and its concentration was measured by a NanoDrop Spectrophotometer (ThermoFisher Scientific). Utilized primers are listed in Table S3.

### E-PAP treatment and measurements of U- and A-tailed RNA frequencies

To measure U-tailed RNA frequencies, 4 µg of TURBO DNase (Invitrogen) treated total RNA was mixed with 0.1 ng *in vitro* transcribed spike-in RNA mixture. This RNA cocktail was treated with *E. coli* poly(A) polymerase (E-PAP, Invitrogen) at 37°C for 30 min in 50 µL reactions, containing 1× reaction buffer, 2.5 mM MnCl_2_, 1 U poly(A) polymerase, 2 U RiboLock RNase Inhibitor (ThermoFisher Scientific) and 1 mM ATP. E-PAP treated RNA was purified using PureLink micro RNA purification kit (Invitrogen) following the manufacturer’s instructions. For the selective RT of U-tailed RNA 10 pmol of the U-tailed RNA specific primer dT_20-_A_4_ primer was used in a 20 µl reaction at 58D to achieve high annealing specificity. RT of all RNA isoforms was carried out with 20 pmol of the dT_20-_ VN primer in a 20 µl reaction at normal PCR extension temperature of 50D. qPCR was carried separately from cDNA synthesized with dT_20-_ A_4_ and dT_20-_ VN primers. U-tailed RNA frequencies were calculated by dividing levels of U-tailed RNA by levels of all isoform RNA. To identify the optimal U-tailed RNA specific primer, five distinct RT primers, dT_20-_A_1_, dT_20-_A_2_, dT_20-_A_3_, dT_20-_A_4_, and dT_20-_A_5_ were tested using the *in vitro* transcribed spike-in RNAs. The dT_20-_A_4_ primer produced minimal background and was chosen for experiments.

To measure A-tailed RNA frequencies, RNA was treated with or without E-PAP as above. RT was carried out with the dT_20-_ VN primer for subsequent qPCR analysis. All isoform levels were measured from samples with E-PAP treatment and A-tailed RNA levels were measured from samples without E-PAP treatment. A-tailed RNA frequencies were calculated by dividing levels of A-tailed RNAs to those of all isoform RNAs.

### RNA immunoprecipitation (RIP)

HeLa cells expressing ‘localization and affinity purification’ (LAP) control or PHAX-LAP constructs ^70,71^ were treated with relevant siRNAs for 3 days. Cells were lysed in extraction buffer (20 mM Tris-HCl pH7.4, 150 mM NaCl, 2 mM EDTA, 1 mM DTT, 1 mM PMSF, 0.1% TritonX-100, supplemented with 1x proteinase inhibitor and 200 U/ml RiboLock RNase Inhibitor) and sonicated using a microtip sonicator (Branson 250). After sonication, lysates were centrifuged, and clarified supernatants were incubated with Dynabeads Epoxy M270 (Invitrogen) conjugated with anti-GFP antibody and rotated at 4LJ overnight. Beads were subsequently washed three times with extraction buffer. 10% volumes of beads were utilized to elute the captured protein using LDS Sample Buffer (Invitrogen) supplemented with Sample Reducing Agent (Invitrogen) for western blotting analysis. The remaining bead volumes were used for RNA extraction using TRIzol (Invitrogen).

### Puromycin incorporation assay

HeLa cells were transfected with siRNAs for 48 hr. To avoid overconfluent cells, resulting in less active translation, cells were reseeded at low confluency (reaching around 50% of confluency the next day) and cultured for another 24 hr. 5 ug/ml of puromycin (Gibco) was added to cell culture medium for 30 min before cell harvest and western blotting analysis.

### Flow cytometry analysis

siRNA transfected HeLa cells were fixed with 70% of cold ethanol for 30 min on ice. Cells were washed with PBS and incubated in a solution, containing 50 ug/ml of propidium iodide (PI, Sigma) and 20 ug/ml of RNase A (ThermoFisher Scientific) at 37LJ for 30 min. Cells were subsequently analyzed in a CytoFLEX machine (Beckman).

### RNA 3’end seq processing and read counts generation

Processing of raw reads and mapping to the genome was done as described (Wu et al., 2020) and mapped reads were then sorted according to their type of non-aligned tail using a custom python script. Reads were stratified via the cigar string information from bam files. Non-templated tail sequences were placed at the 5’end of reads in the utilized library preparation method (Lexogen QuantSeq REV) and thus identified with cigar strings matching the regular expressions “^.S..M” and “^..S..M” for positive- and “..M.S$” or “..M..S$” for negative-strand mappings. Further downstream analysis of total tail counts, tail-specific filtering and the generation of coverage tracks were performed using custom python and bash scripts.

### Frequency plots

Boxplots displaying the frequencies of U-tails within samples, the representation of abundance of the individual tails’ motives and the display of U-containing reads relative to tail length within genomic and spike-in RNA were all based on counts mentioned above and generated using the ggplot2 R package.

### Metagene profiles and heatmaps

Metagene profiles and heatmaps were produced using custom R scripts as in (Lykke-Andersen et al., 2021). Briefly, the rtracklayer R package was used to collect read coverage values for the window -1kb/+6kb according to the TSS. Coverage values were then binned in 50nt bins and used to generate heatmaps based on the R package ComplexHeatmap. The mean of coverage values across TUs over each bin were also computed and plotted as metagene profiles using custom R code. A 95% confidence interval of the mean coverage is displayed for each sample and was measured through 50 steps of bootstrap samplings with replacement.

### Genome Browser views

Genome browser views were generated using BigWig files and the R package seqNdisplayR (https://rdrr.io/github/THJlab/seqNdisplayR/).

---

**Figure S1, related to Figure 1.**
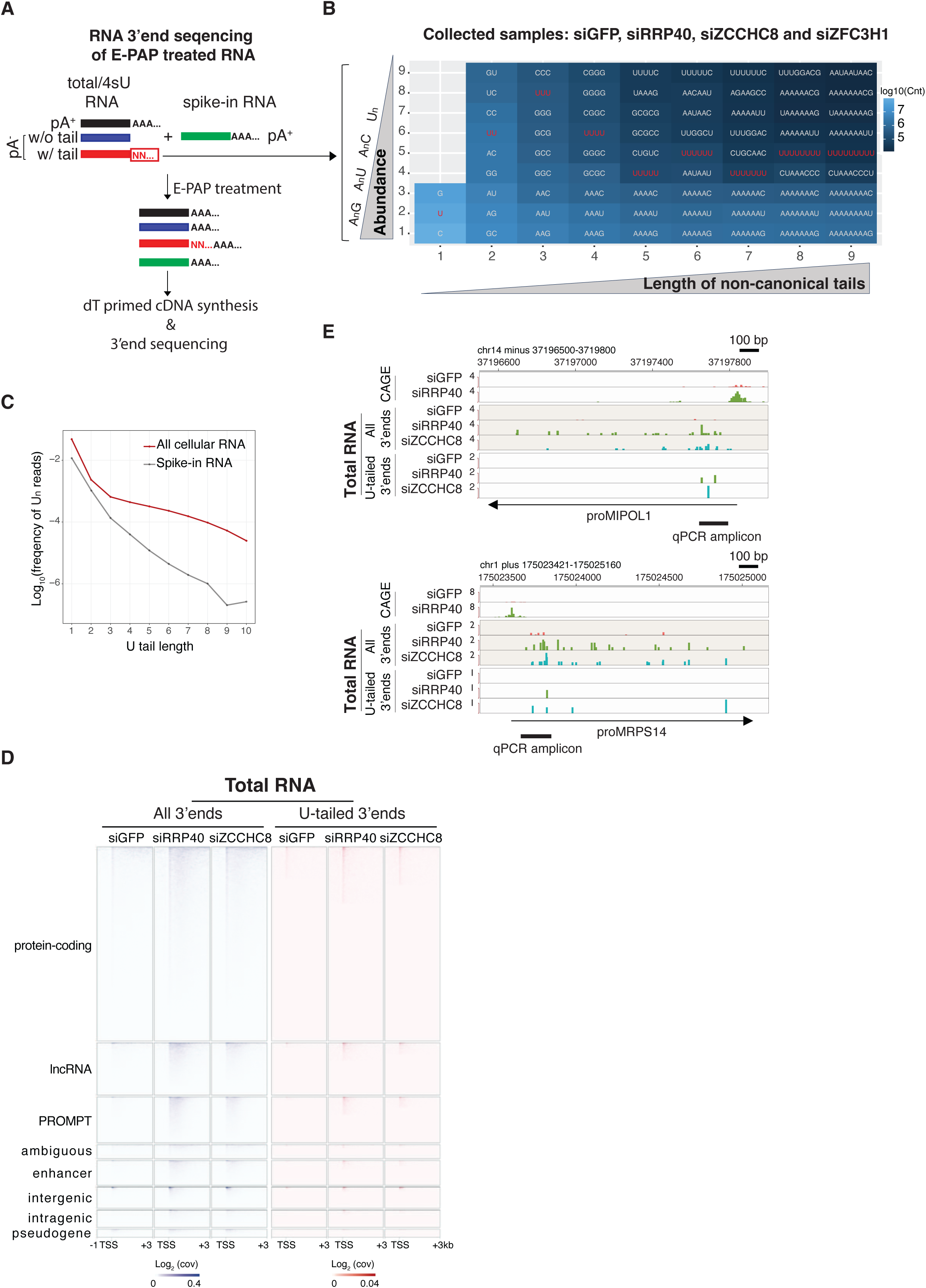
**(A)** Workflow of the RNA 3’end seq approach. Total- and 4sU-RNA were mixed with spike-in RNAs and treated with *E.coli* poly(A) polymerase (E-PAP). cDNA synthesis was conducted with an oligo dT primer prior to library preparation and 3’end sequencing. The red box and black arrow denote the non-canonical tails presented in Figure S1B after bioinformatics analysis. NN = at least one non-adenosine nucleotide. **(B)** Quantification of non-canonical tails. The visualized data were combined from E-PAP-treated RNAs from the indicated depletion conditions with each sample providing biological replicates of total- (3) and 4sU- (2) RNA (GEO: GSE137612, ^36^). Tails harboring from 1 to 9 nt (x-axis) were sorted according to their abundance (y-axis). Tails containing adenosine nucleotides only were excluded from analysis. **(C)** Quantification of U-tail frequencies in cellular *vs.* spike-in RNA. U-tails from 1-10 nt in length are shown. Data from Figure S1B were used. **(D)** Heatmaps as in Figure 1C, but showing total RNA data. **(E)** Genome browser views as in Figure 1E, but showing total RNA data.

**Figure S2, related to Figure 2.**
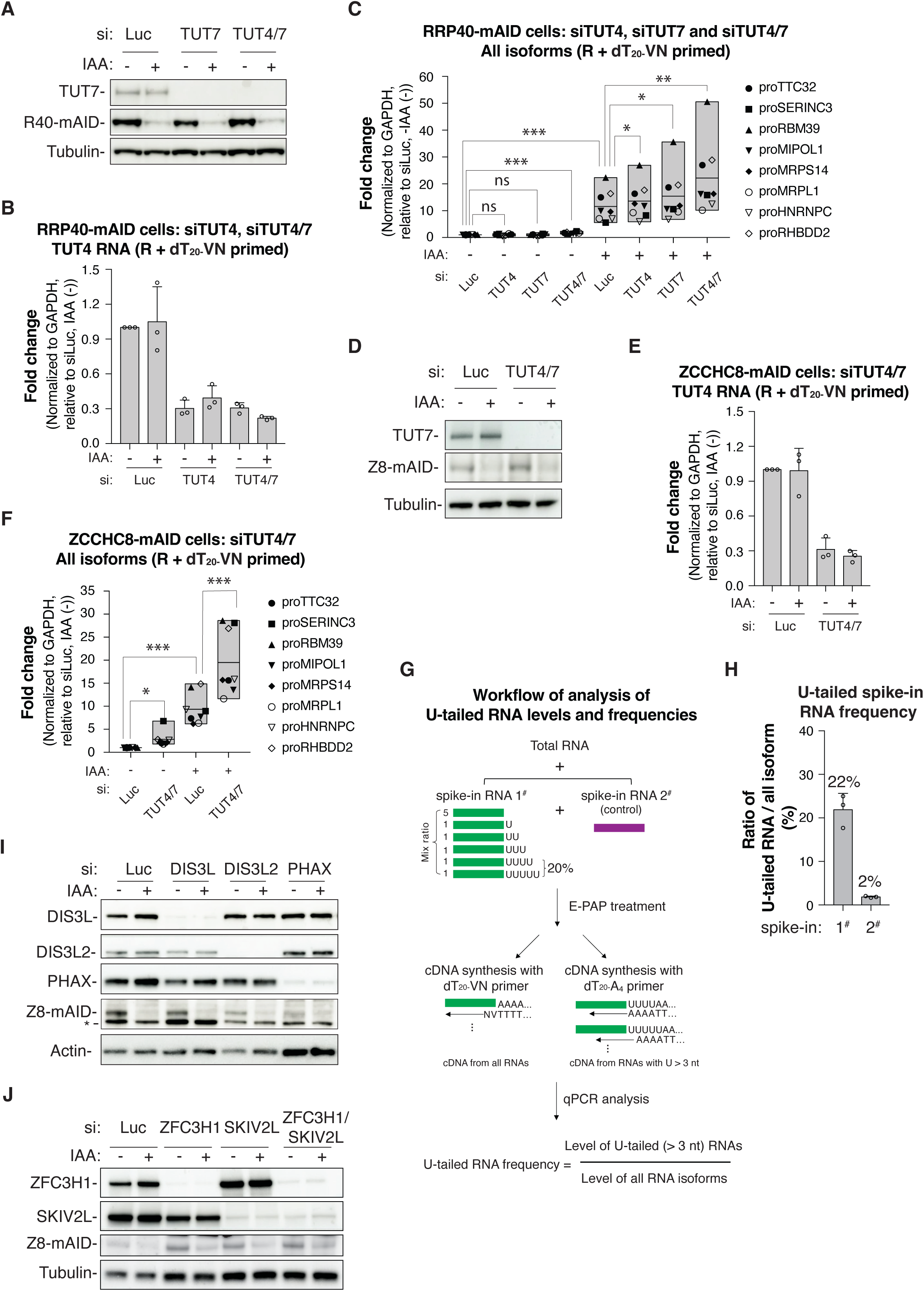
**(A)** Western blotting analysis of the indicated IAA- and siRNA-mediated protein depletions employing RRP40-mAID cells. Luciferase (Luc) siRNA was used as a negative control. Western membranes were probed with antibodies towards RRP40-mAID (R40-mAID), TUT7 and Tubulin as a loading control. **(B)** Bar plots of qRT-PCR analysis of TUT4 RNA levels from samples as indicated. Biological triplicate data are shown with each circle representing single replicate values. qRT-PCR data were normalized to GAPDH RNA levels and relative to the control sample (siLuc, IAA (-)). **(C)** Boxplots of qRT-PCR analysis as in Figure 2A, but employing siLuc, siTUT4 and/or siTUT7 siRNAs as indicated. Moreover, cDNA was synthesized using a mix of random hexamer- and dT_20-_VN-primers (R+dT_20-_VN). Data normalization and statistical analysis as in Figure 2A. Note that boxplots, which display measurements of all RNA isoform levels after R+dT_20-_VN priming condition, are consistently represented in gray color. **(D)** Western blotting analysis as in (A), but employing ZCCHC8-mAID cells and a combination of siTUT4 and siTUT7. Western membranes were probed with antibodies towards TUT7, ZCCHC8-mAID (Z8-mAID), and Tubulin as a loading control. **(E)** Bar plots of qRT-PCR analysis as in (B), but using samples from (D). **(F)** Boxplots of qRT-PCR analysis as in (C), but employing ZCCHC8-mAID cells. **(G)** Workflow of analysis to estimate U-tailed RNA frequencies. Total cellular RNA was mixed with spike-in RNA 1^#^ and 2^#^ species as indicated. After E-PAP treatment, cDNA was produced, using dT_20_-NV or dT_20_A_4_ primers, respectively, enabling measurements of all RNA isoform levels and consequent calculation of U-tailed RNA frequencies. **(H)** Measurement of U-tail frequency of spike-in RNA (as outlined in (G)). Biological triplicate data are shown with each circle representing one replicate. Measured frequencies are shown for each spike-in RNA sample. **(I)** Western blotting analysis of the indicated IAA- and siRNA-mediated protein depletions employing ZCCHC8-mAID cells. Western membranes were probed with antibodies towards DIS3L, DIS3L2, PHAX (used later), ZCCHC8-mAID (Z8-mAID) and Actin as a loading control. *Denotes a non-specific band. **(J)** Western blotting analysis of the indicated IAA- and siRNA-mediated protein depletions, employing ZCCHC8-mAID cells. Western membranes were probed with antibodies towards ZFC3H1 (used later), SKIV2L, ZCCHC8-mAID (Z8-mAID) and Tubulin as a loading control.

**Figure S3, related to Figure 3.**
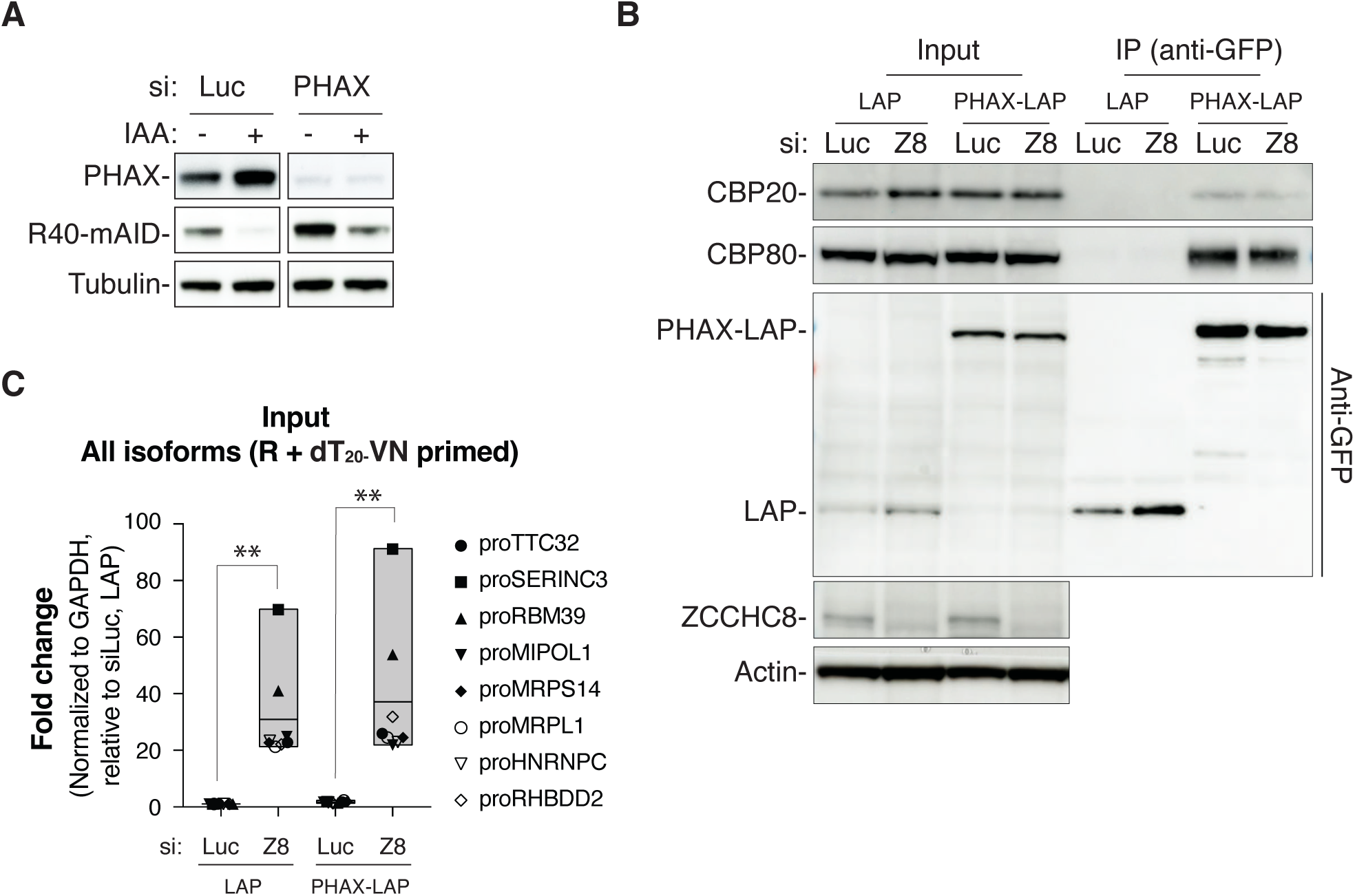
**(A)** Western blotting analysis of the indicated IAA- and siRNA-mediated protein depletions, employing RRP40-mAID cells. Western membranes were probed with antibodies towards PHAX, RRP40-mAID (R40-mAID) and Tubulin as a loading control. **(B)** Western blotting analysis of the indicated proteins co-purifying with LAP (negative control) or PHAX-LAP, employing a GFP antibody-IP from extracts of cells transfected with siLuc or siZCCHC8 (siZ8) as indicated. Input and IP samples were probed with antibodies towards CBP20, CBP80, and GFP. Membrane with input samples was further probed with antibodies towards ZCCHC8 and Actin (loading control) to confirm ZCCHC8 depletion. **(C)** Boxplots of qRT-PCR analysis as in Figure 3E, but employing the input samples from Figure S3B.

**Figure S4, related to Figure 4.**
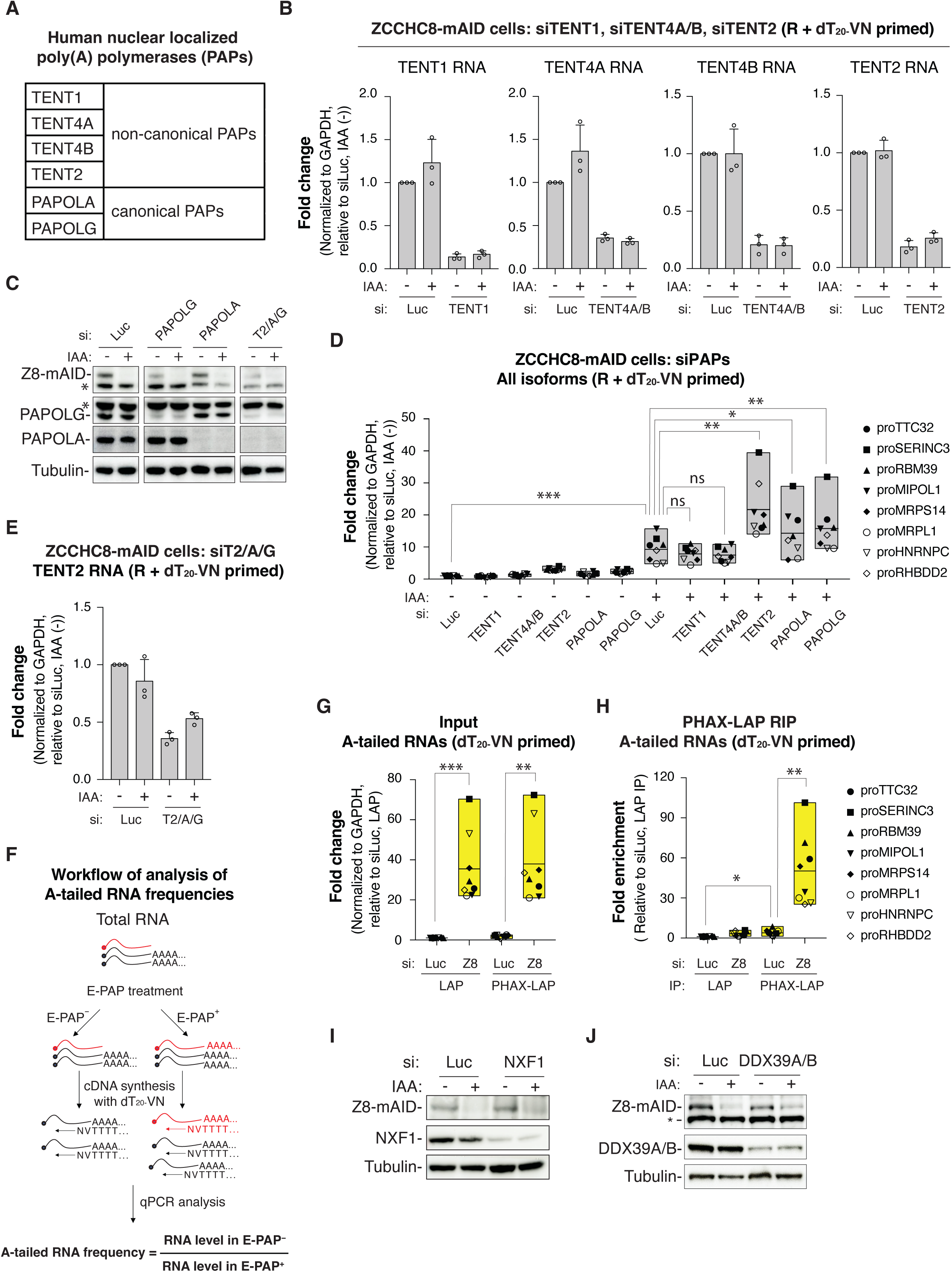
**(A)** Table of nuclear-localized PAPs expressed in HeLa cells. **(B)** Bar plots of qRT-PCR analysis as in Figure S2B, but analyzing TENT1, TENT4A, TENT4B and TENT2 RNA levels in the indicated IAA- and siRNA-mediated depletion samples from ZCCHC8-mAID cells. **(C)** Western blotting analysis of the indicated IAA- and siRNA-mediated protein depletions, employing ZCCHC8-mAID cells. Western membranes were probed with antibodies towards ZCCHC8-mAID (Z8-mAID), PAPOLG, PAPOLA and Tubulin as a loading control. *Denotes non-specific bands. siT2/A/G (siTENT2/PAPOLA/PAPOLG) samples were used later. **(D)** Boxplots of qRT-PCR analysis as in Figure S2F, but employing siLuc, siTENT1, siTENT4A/B, siTENT2, siPAPOLA and siPAPOLG siRNAs as indicated. **(E)** Bar plots of qRT-PCR analysis as in Figure S2B, but analyzing TENT2 RNA levels in siLuc and siT2/A/G (siTENT2/PAPOLA/PAPOLG) samples from (C). **(F)** Workflow of analysis of A-tailed RNA frequencies. Total RNA was treated with (+) or without (-) E-PAP and reverse transcribed using a dT_20-_VN primer before qPCR analysis. A-tailed RNA frequencies were calculated by dividing RNA levels in E-PAP^-^ samples to those in E-PAP^+^ samples. **(G)** Boxplots of qRT-PCR analysis as in Figure S3C, but synthesizing cDNA using a dT_20-_VN primer. **(H)** Boxplots of qRT-PCR analysis as (F), but employing IP samples. **(I)** Western blotting analysis of the indicated IAA- and siRNA-mediated protein depletions, employing ZCCHC8-mAID cells. Western membranes were probed with antibodies towards ZCCHC8-mAID (Z8-mAID), NXF1 and Tubulin as a loading control. **(J)** Western blotting analysis as in (I), but depleting DDX39A/B. *Denotes a non-specific band.

**Figure S5, related to Figure 5.**
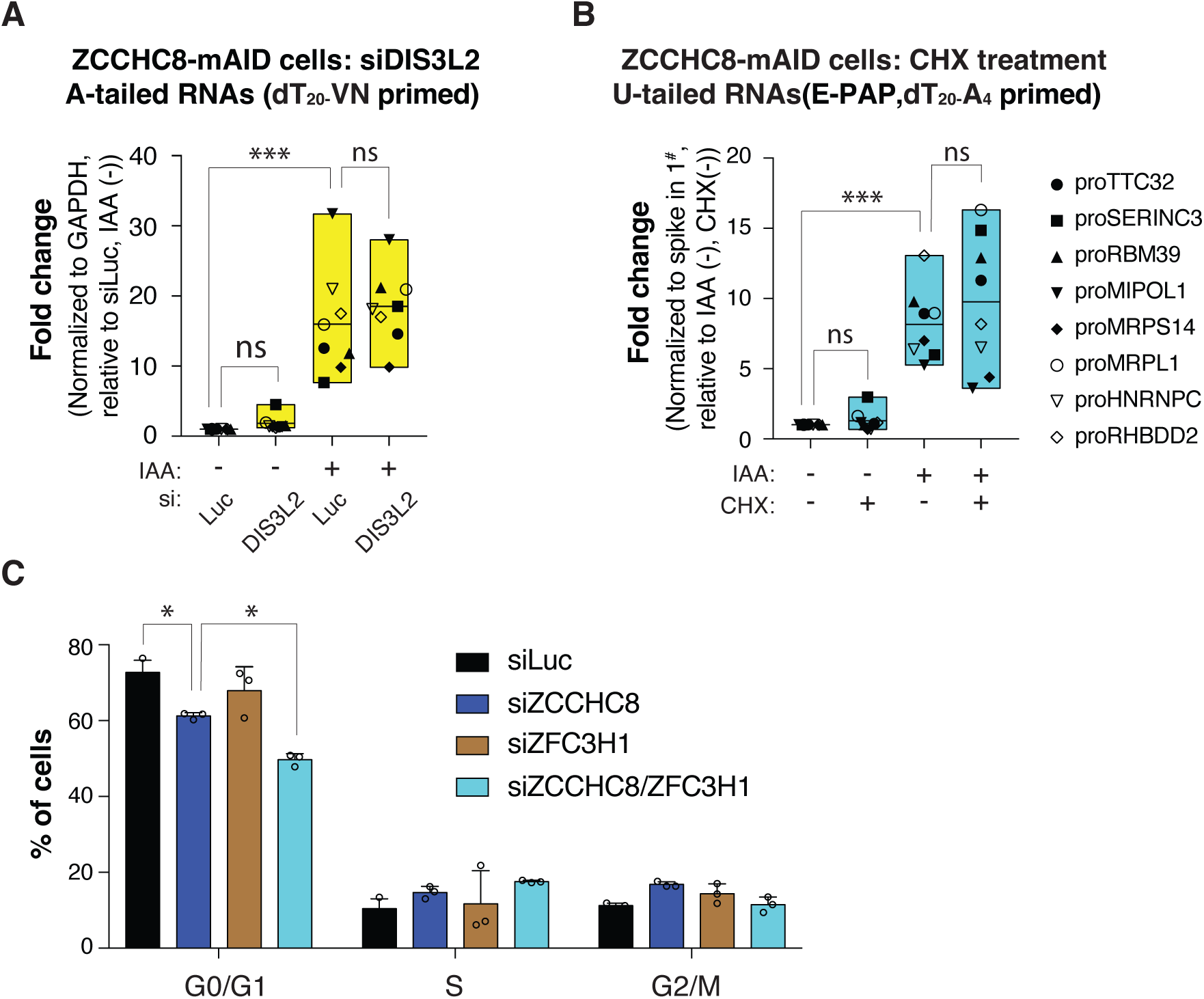
**(A)** Boxplots of qRT-PCR analysis as in Figure 5A, but employing the DIS3L2 depletion samples from Figure S2I. **(B)** Boxplots of qRT-PCR analysis as in Figure 5C, but measuring U-tailed RNA levels. Note that this box plot, which displays measurement of U-tailed RNA levels, is represented in blue color. **(C)** Quantification G0/G1, S and G2/M phase cell population data from Figure 5F. Biological triplicate data are shown with each circle representing single replicate values.

**Figure S6, related to Figure 6.**
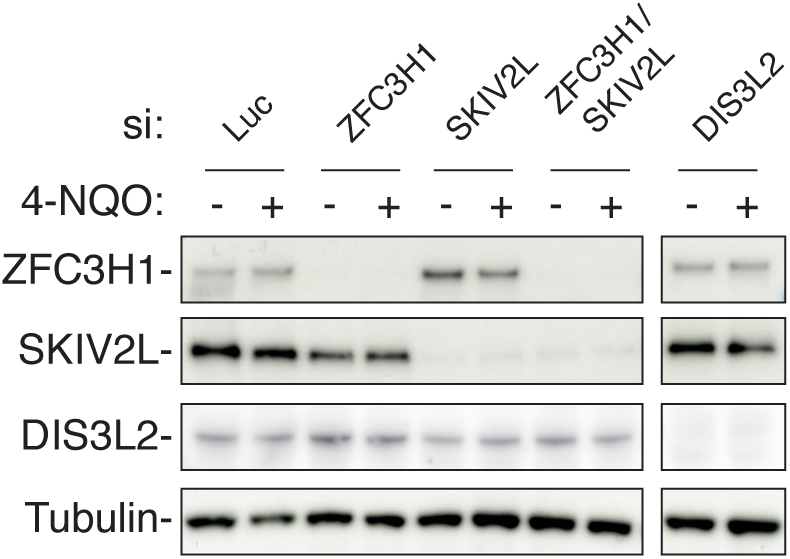
Western blotting analysis of the indicated siRNA-mediated protein depletion samples, co-treated with (+) or without (-) 4-NQO. Western membranes were probed with antibodies towards ZFC3H1, SKIV2L, DIS3L2 and Tubulin as a loading control.

## References

1. Jensen, T.H., Jacquier, A., and Libri, D. (2013). Dealing with pervasive transcription. Mol Cell 52, 473–484. 10.1016/j.molcel.2013.10.032.

2. Jacquier, A. (2009). The complex eukaryotic transcriptome: unexpected pervasive transcription and novel small RNAs. Nat Rev Genet 10, 833–844. 10.1038/nrg2683.

3. Doma, M.K., and Parker, R. (2007). RNA quality control in eukaryotes. Cell 131, 660–668. 10.1016/j.cell.2007.10.041.

4. Schmid, M., and Jensen, T.H. (2018). Controlling nuclear RNA levels. Nat Rev Genet 19, 518–529. 10.1038/s41576-018-0013-2.

5. Staals, R.H., Bronkhorst, A.W., Schilders, G., Slomovic, S., Schuster, G., Heck, A.J., Raijmakers, R., and Pruijn, G.J. (2010). Dis3-like 1: a novel exoribonuclease associated with the human exosome. EMBO J 29, 2358–2367. 10.1038/emboj.2010.122.

6. Tomecki, R., Kristiansen, M.S., Lykke-Andersen, S., Chlebowski, A., Larsen, K.M., Szczesny, R.J., Drazkowska, K., Pastula, A., Andersen, J.S., Stepien, P.P., et al. (2010). The human core exosome interacts with differentially localized processive RNases: hDIS3 and hDIS3L. EMBO J 29, 2342–2357. 10.1038/emboj.2010.121.

7. Anderson, J.S., and Parker, R.P. (1998). The 3’ to 5’ degradation of yeast mRNAs is a general mechanism for mRNA turnover that requires the SKI2 DEVH box protein and 3’ to 5’ exonucleases of the exosome complex. EMBO J 17, 1497–1506. 10.1093/emboj/17.5.1497.

8. Halbach, F., Reichelt, P., Rode, M., and Conti, E. (2013). The yeast ski complex: crystal structure and RNA channeling to the exosome complex. Cell 154, 814–826. 10.1016/j.cell.2013.07.017.

9. Kogel, A., Keidel, A., Bonneau, F., Schafer, I.B., and Conti, E. (2022). The human SKI complex regulates channeling of ribosome-bound RNA to the exosome via an intrinsic gatekeeping mechanism. Mol Cell 82, 756–769 e758. 10.1016/j.molcel.2022.01.009.

10. Wiederhold, K., and Passmore, L.A. (2010). Cytoplasmic deadenylation: regulation of mRNA fate. Biochem Soc Trans 38, 1531–1536. 10.1042/BST0381531.

11. Eisen, T.J., Eichhorn, S.W., Subtelny, A.O., Lin, K.S., McGeary, S.E., Gupta, S., and Bartel, D.P. (2020). The Dynamics of Cytoplasmic mRNA Metabolism. Mol Cell 77, 786–799 e710. 10.1016/j.molcel.2019.12.005.

12. Malecki, M., Viegas, S.C., Carneiro, T., Golik, P., Dressaire, C., Ferreira, M.G., and Arraiano, C.M. (2013). The exoribonuclease Dis3L2 defines a novel eukaryotic RNA degradation pathway. EMBO J 32, 1842–1854. 10.1038/emboj.2013.63.

13. Chang, H.M., Triboulet, R., Thornton, J.E., and Gregory, R.I. (2013). A role for the Perlman syndrome exonuclease Dis3l2 in the Lin28-let-7 pathway. Nature 497, 244–248. 10.1038/nature12119.

14. Lubas, M., Damgaard, C.K., Tomecki, R., Cysewski, D., Jensen, T.H., and Dziembowski, A. (2013). Exonuclease hDIS3L2 specifies an exosome-independent 3’-5’ degradation pathway of human cytoplasmic mRNA. EMBO J 32, 1855–1868. 10.1038/emboj.2013.135.

15. Faehnle, C.R., Walleshauser, J., and Joshua-Tor, L. (2014). Mechanism of Dis3l2 substrate recognition in the Lin28-let-7 pathway. Nature 514, 252–256. 10.1038/nature13553.

16. De Almeida, C., Scheer, H., Zuber, H., and Gagliardi, D. (2018). RNA uridylation: a key posttranscriptional modification shaping the coding and noncoding transcriptome. Wiley Interdiscip Rev RNA 9. 10.1002/wrna.1440.

17. Ustianenko, D., Pasulka, J., Feketova, Z., Bednarik, L., Zigackova, D., Fortova, A., Zavolan, M., and Vanacova, S. (2016). TUT-DIS3L2 is a mammalian surveillance pathway for aberrant structured non-coding RNAs. EMBO J 35, 2179–2191. 10.15252/embj.201694857.

18. Okamura, M., Inose, H., and Masuda, S. (2015). RNA Export through the NPC in Eukaryotes. Genes (Basel) 6, 124–149. 10.3390/genes6010124.

19. Kohler, A., and Hurt, E. (2007). Exporting RNA from the nucleus to the cytoplasm. Nat Rev Mol Cell Biol 8, 761–773. 10.1038/nrm2255.

20. McCloskey, A., Taniguchi, I., Shinmyozu, K., and Ohno, M. (2012). hnRNP C tetramer measures RNA length to classify RNA polymerase II transcripts for export. Science 335, 1643–1646. 10.1126/science.1218469.

21. Ohno, M., Segref, A., Bachi, A., Wilm, M., and Mattaj, I.W. (2000). PHAX, a mediator of U snRNA nuclear export whose activity is regulated by phosphorylation. Cell 101, 187–198. 10.1016/S0092-8674(00)80829-6.

22. Masuyama, K., Taniguchi, I., Kataoka, N., and Ohno, M. (2004). RNA length defines RNA export pathway. Genes Dev 18, 2074–2085. 10.1101/gad.1216204.

23. Machitani, M., Taniguchi, I., McCloskey, A., Suzuki, T., and Ohno, M. (2020). The RNA transport factor PHAX is required for proper histone H2AX expression and DNA damage response. RNA 26, 1716–1725. 10.1261/rna.074625.120.

24. Viphakone, N., Hautbergue, G.M., Walsh, M., Chang, C.T., Holland, A., Folco, E.G., Reed, R., and Wilson, S.A. (2012). TREX exposes the RNA-binding domain of Nxf1 to enable mRNA export. Nat Commun 3, 1006. 10.1038/ncomms2005.

25. Carmody, S.R., and Wente, S.R. (2009). mRNA nuclear export at a glance. J Cell Sci 122, 1933–1937. 10.1242/jcs.041236.

26. Mitchell, P., Petfalski, E., Shevchenko, A., Mann, M., and Tollervey, D. (1997). The exosome: a conserved eukaryotic RNA processing complex containing multiple 3’-->5’ exoribonucleases. Cell 91, 457–466. 10.1016/s0092-8674(00)80432-8.

27. Wasmuth, E.V., Zinder, J.C., Zattas, D., Das, M., and Lima, C.D. (2017). Structure and reconstitution of yeast Mpp6-nuclear exosome complexes reveals that Mpp6 stimulates RNA decay and recruits the Mtr4 helicase. Elife 6. 10.7554/eLife.29062.

28. Falk, S., Bonneau, F., Ebert, J., Kogel, A., and Conti, E. (2017). Mpp6 Incorporation in the Nuclear Exosome Contributes to RNA Channeling through the Mtr4 Helicase. Cell Rep 20, 2279–2286. 10.1016/j.celrep.2017.08.033.

29. Schuch, B., Feigenbutz, M., Makino, D.L., Falk, S., Basquin, C., Mitchell, P., and Conti, E. (2014). The exosome-binding factors Rrp6 and Rrp47 form a composite surface for recruiting the Mtr4 helicase. EMBO J 33, 2829–2846. 10.15252/embj.201488757.

30. Lubas, M., Christensen, M.S., Kristiansen, M.S., Domanski, M., Falkenby, L.G., Lykke-Andersen, S., Andersen, J.S., Dziembowski, A., and Jensen, T.H. (2011). Interaction profiling identifies the human nuclear exosome targeting complex. Mol Cell 43, 624–637. 10.1016/j.molcel.2011.06.028.

31. Meola, N., Domanski, M., Karadoulama, E., Chen, Y., Gentil, C., Pultz, D., Vitting-Seerup, K., Lykke-Andersen, S., Andersen, J.S., Sandelin, A., and Jensen, T.H. (2016). Identification of a Nuclear Exosome Decay Pathway for Processed Transcripts. Mol Cell 64, 520–533. 10.1016/j.molcel.2016.09.025.

32. Silla, T., Schmid, M., Dou, Y., Garland, W., Milek, M., Imami, K., Johnsen, D., Polak, P., Andersen, J.S., Selbach, M., et al. (2020). The human ZC3H3 and RBM26/27 proteins are critical for PAXT-mediated nuclear RNA decay. Nucleic Acids Res 48, 2518–2530. 10.1093/nar/gkz1238.

33. Gerlach, P., Garland, W., Lingaraju, M., Salerno-Kochan, A., Bonneau, F., Basquin, J., Jensen, T.H., and Conti, E. (2022). Structure and regulation of the nuclear exosome targeting complex guides RNA substrates to the exosome. Mol Cell 82, 2505–2518 e2507. 10.1016/j.molcel.2022.04.011.

34. Puno, M.R., and Lima, C.D. (2022). Structural basis for RNA surveillance by the human nuclear exosome targeting (NEXT) complex. Cell 185, 2132–2147 e2126. 10.1016/j.cell.2022.04.016.

35. Andersen, P.R., Domanski, M., Kristiansen, M.S., Storvall, H., Ntini, E., Verheggen, C., Schein, A., Bunkenborg, J., Poser, I., Hallais, M., et al. (2013). The human cap-binding complex is functionally connected to the nuclear RNA exosome. Nat Struct Mol Biol 20, 1367–1376. 10.1038/nsmb.2703.

36. Wu, G., Schmid, M., Rib, L., Polak, P., Meola, N., Sandelin, A., and Jensen, T.H. (2020). A Two-Layered Targeting Mechanism Underlies Nuclear RNA Sorting by the Human Exosome. Cell Rep 30, 2387–2401 e2385. 10.1016/j.celrep.2020.01.068.

37. Dubiez, E., Pellegrini, E., Finderup Brask, M., Garland, W., Foucher, A.E., Huard, K., Heick Jensen, T., Cusack, S., and Kadlec, J. (2024). Structural basis for competitive binding of productive and degradative co-transcriptional effectors to the nuclear cap-binding complex. Cell Rep 43, 113639. 10.1016/j.celrep.2023.113639.

38. Ogami, K., Richard, P., Chen, Y., Hoque, M., Li, W., Moresco, J.J., Yates, J.R., 3rd, Tian, B., and Manley, J.L. (2017). An Mtr4/ZFC3H1 complex facilitates turnover of unstable nuclear RNAs to prevent their cytoplasmic transport and global translational repression. Genes Dev 31, 1257–1271. 10.1101/gad.302604.117.

39. Bresson, S.M., Hunter, O.V., Hunter, A.C., and Conrad, N.K. (2015). Canonical Poly(A) Polymerase Activity Promotes the Decay of a Wide Variety of Mammalian Nuclear RNAs. PLoS Genet 11, e1005610. 10.1371/journal.pgen.1005610.

40. Polak, P., Garland, W., Rathore, O., Schmid, M., Salerno-Kochan, A., Jakobsen, L., Gockert, M., Gerlach, P., Silla, T., Andersen, J.S., et al. (2023). Dual agonistic and antagonistic roles of ZC3H18 provide for co-activation of distinct nuclear RNA decay pathways. Cell Rep 42, 113325. 10.1016/j.celrep.2023.113325.

41. Bugai, A., Quaresma, A.J.C., Friedel, C.C., Lenasi, T., Duster, R., Sibley, C.R., Fujinaga, K., Kukanja, P., Hennig, T., Blasius, M., et al. (2019). P-TEFb Activation by RBM7 Shapes a Pro-survival Transcriptional Response to Genotoxic Stress. Mol Cell 74, 254–267 e210. 10.1016/j.molcel.2019.01.033.

42. Tiedje, C., Lubas, M., Tehrani, M., Menon, M.B., Ronkina, N., Rousseau, S., Cohen, P., Kotlyarov, A., and Gaestel, M. (2015). p38MAPK/MK2-mediated phosphorylation of RBM7 regulates the human nuclear exosome targeting complex. RNA 21, 262–278. 10.1261/rna.048090.114.

43. Blasius, M., Wagner, S.A., Choudhary, C., Bartek, J., and Jackson, S.P. (2014). A quantitative 14-3-3 interaction screen connects the nuclear exosome targeting complex to the DNA damage response. Genes Dev 28, 1977–1982. 10.1101/gad.246272.114.

44. Nechaev, S., Fargo, D.C., dos Santos, G., Liu, L., Gao, Y., and Adelman, K. (2010). Global analysis of short RNAs reveals widespread promoter-proximal stalling and arrest of Pol II in Drosophila. Science 327, 335–338. 10.1126/science.1181421.

45. Kamieniarz-Gdula, K., and Proudfoot, N.J. (2019). Transcriptional Control by Premature Termination: A Forgotten Mechanism. Trends Genet 35, 553–564. 10.1016/j.tig.2019.05.005.

46. Tatomer, D.C., and Wilusz, J.E. (2019). Attenuation of Eukaryotic Protein-Coding Gene Expression via Premature Transcription Termination. Cold Spring Harb Symp Quant Biol 84, 83–93. 10.1101/sqb.2019.84.039644.

47. Elrod, N.D., Henriques, T., Huang, K.L., Tatomer, D.C., Wilusz, J.E., Wagner, E.J., and Adelman, K. (2019). The Integrator Complex Attenuates Promoter-Proximal Transcription at Protein-Coding Genes. Mol Cell 76, 738–752 e737. 10.1016/j.molcel.2019.10.034.

48. Tatomer, D.C., Elrod, N.D., Liang, D., Xiao, M.S., Jiang, J.Z., Jonathan, M., Huang, K.L., Wagner, E.J., Cherry, S., and Wilusz, J.E. (2019). The Integrator complex cleaves nascent mRNAs to attenuate transcription. Genes Dev 33, 1525–1538. 10.1101/gad.330167.119.

49. Beckedorff, F., Blumenthal, E., daSilva, L.F., Aoi, Y., Cingaram, P.R., Yue, J., Zhang, A., Dokaneheifard, S., Valencia, M.G., Gaidosh, G., et al. (2020). The Human Integrator Complex Facilitates Transcriptional Elongation by Endonucleolytic Cleavage of Nascent Transcripts. Cell Rep 32, 107917. 10.1016/j.celrep.2020.107917.

50. Lykke-Andersen, S., Zumer, K., Molska, E.S., Rouviere, J.O., Wu, G., Demel, C., Schwalb, B., Schmid, M., Cramer, P., and Jensen, T.H. (2021). Integrator is a genome-wide attenuator of non-productive transcription. Mol Cell 81, 514–529 e516. 10.1016/j.molcel.2020.12.014.

51. Estell, C., Davidson, L., Eaton, J.D., Kimura, H., Gold, V.A.M., and West, S. (2023). A restrictor complex of ZC3H4, WDR82, and ARS2 integrates with PNUTS to control unproductive transcription. Mol Cell 83, 2222–2239 e2225. 10.1016/j.molcel.2023.05.029.

52. Rouviere, J.O., Salerno-Kochan, A., Lykke-Andersen, S., Garland, W., Dou, Y., Rathore, O., Molska, E.S., Wu, G., Schmid, M., Bugai, A., et al. (2023). ARS2 instructs early transcription termination-coupled RNA decay by recruiting ZC3H4 to nascent transcripts. Mol Cell 83, 2240–2257 e2246. 10.1016/j.molcel.2023.05.028.

53. Lim, J., Kim, D., Lee, Y.S., Ha, M., Lee, M., Yeo, J., Chang, H., Song, J., Ahn, K., and Kim, V.N. (2018). Mixed tailing by TENT4A and TENT4B shields mRNA from rapid deadenylation. Science 361, 701–704. 10.1126/science.aam5794.

54. Warkocki, Z., Krawczyk, P.S., Adamska, D., Bijata, K., Garcia-Perez, J.L., and Dziembowski, A. (2018). Uridylation by TUT4/7 Restricts Retrotransposition of Human LINE-1s. Cell 174, 1537–1548 e1529. 10.1016/j.cell.2018.07.022.

55. Kim, D., Lee, Y.S., Jung, S.J., Yeo, J., Seo, J.J., Lee, Y.Y., Lim, J., Chang, H., Song, J., Yang, J., et al. (2020). Viral hijacking of the TENT4-ZCCHC14 complex protects viral RNAs via mixed tailing. Nat Struct Mol Biol 27, 581–588. 10.1038/s41594-020-0427-3.

56. Hyrina, A., Jones, C., Chen, D., Clarkson, S., Cochran, N., Feucht, P., Hoffman, G., Lindeman, A., Russ, C., Sigoillot, F., et al. (2019). A Genome-wide CRISPR Screen Identifies ZCCHC14 as a Host Factor Required for Hepatitis B Surface Antigen Production. Cell Rep 29, 2970–2978 e2976. 10.1016/j.celrep.2019.10.113.

57. Yashiro, Y., and Tomita, K. (2018). Function and Regulation of Human Terminal Uridylyltransferases. Front Genet 9, 538. 10.3389/fgene.2018.00538.

58. Yu, S., and Kim, V.N. (2020). A tale of non-canonical tails: gene regulation by post-transcriptional RNA tailing. Nat Rev Mol Cell Biol 21, 542–556. 10.1038/s41580-020-0246-8.

59. Liudkovska, V., and Dziembowski, A. (2021). Functions and mechanisms of RNA tailing by metazoan terminal nucleotidyltransferases. Wiley Interdiscip Rev RNA 12, e1622. 10.1002/wrna.1622.

60. Ma, X., Shao, Y., Tian, L., Flasch, D.A., Mulder, H.L., Edmonson, M.N., Liu, Y., Chen, X., Newman, S., Nakitandwe, J., et al. (2019). Analysis of error profiles in deep next-generation sequencing data. Genome Biol 20, 50. 10.1186/s13059-019-1659-6.

61. Stoler, N., and Nekrutenko, A. (2021). Sequencing error profiles of Illumina sequencing instruments. NAR Genom Bioinform 3, lqab019. 10.1093/nargab/lqab019.

62. Warkocki, Z., Liudkovska, V., Gewartowska, O., Mroczek, S., and Dziembowski, A. (2018). Terminal nucleotidyl transferases (TENTs) in mammalian RNA metabolism. Philos Trans R Soc Lond B Biol Sci 373. 10.1098/rstb.2018.0162.

63. Scheer, H., Zuber, H., De Almeida, C., and Gagliardi, D. (2016). Uridylation Earmarks mRNAs for Degradation… and More. Trends Genet 32, 607–619. 10.1016/j.tig.2016.08.003.

64. Gockert, M., Schmid, M., Jakobsen, L., Jens, M., Andersen, J.S., and Jensen, T.H. (2022). Rapid factor depletion highlights intricacies of nucleoplasmic RNA degradation. Nucleic Acids Res 50, 1583–1600. 10.1093/nar/gkac001.

65. Morgan, M., Kabayama, Y., Much, C., Ivanova, I., Di Giacomo, M., Auchynnikava, T., Monahan, J.M., Vitsios, D.M., Vasiliauskaite, L., Comazzetto, S., et al. (2019). A programmed wave of uridylation-primed mRNA degradation is essential for meiotic progression and mammalian spermatogenesis. Cell Res 29, 221–232. 10.1038/s41422-018-0128-1.

66. Tuck, A.C., Rankova, A., Arpat, A.B., Liechti, L.A., Hess, D., Iesmantavicius, V., Castelo-Szekely, V., Gatfield, D., and Buhler, M. (2020). Mammalian RNA Decay Pathways Are Highly Specialized and Widely Linked to Translation. Mol Cell 77, 1222–1236 e1213. 10.1016/j.molcel.2020.01.007.

67. Meze, K., Axhemi, A., Thomas, D.R., Doymaz, A., and Joshua-Tor, L. (2023). A shape-shifting nuclease unravels structured RNA. Nat Struct Mol Biol 30, 339–347. 10.1038/s41594-023-00923-x.

68. Zigackova, D., and Vanacova, S. (2018). The role of 3’ end uridylation in RNA metabolism and cellular physiology. Philos Trans R Soc Lond B Biol Sci 373. 10.1098/rstb.2018.0171.

69. Kurosaki, T., Miyoshi, K., Myers, J.R., and Maquat, L.E. (2018). NMD-degradome sequencing reveals ribosome-bound intermediates with 3’-end non-templated nucleotides. Nat Struct Mol Biol 25, 940–950. 10.1038/s41594-018-0132-7.

70. Giacometti, S., Benbahouche, N.E.H., Domanski, M., Robert, M.C., Meola, N., Lubas, M., Bukenborg, J., Andersen, J.S., Schulze, W.M., Verheggen, C., et al. (2017). Mutually Exclusive CBC-Containing Complexes Contribute to RNA Fate. Cell Rep 18, 2635–2650. 10.1016/j.celrep.2017.02.046.

71. Poser, I., Sarov, M., Hutchins, J.R., Heriche, J.K., Toyoda, Y., Pozniakovsky, A., Weigl, D., Nitzsche, A., Hegemann, B., Bird, A.W., et al. (2008). BAC TransgeneOmics: a high-throughput method for exploration of protein function in mammals. Nat Methods 5, 409–415. 10.1038/nmeth.1199.

72. De Magistris, P. (2021). The Great Escape: mRNA Export through the Nuclear Pore Complex. Int J Mol Sci 22. 10.3390/ijms222111767.

73. Zinoviev, A., Ayupov, R.K., Abaeva, I.S., Hellen, C.U.T., and Pestova, T.V. (2020). Extraction of mRNA from Stalled Ribosomes by the Ski Complex. Mol Cell 77, 1340–1349 e1346. 10.1016/j.molcel.2020.01.011.

74. Schmidt, E.K., Clavarino, G., Ceppi, M., and Pierre, P. (2009). SUnSET, a nonradioactive method to monitor protein synthesis. Nat Methods 6, 275–277. 10.1038/nmeth.1314.

75. Darken, M.A. (1964). Puromycin Inhibition of Protein Synthesis. Pharmacol Rev 16, 223–243.

76. Nathans, D. (1964). Puromycin Inhibition of Protein Synthesis: Incorporation of Puromycin into Peptide Chains. Proc Natl Acad Sci U S A 51, 585–592. 10.1073/pnas.51.4.585.

77. Malabat, C., Feuerbach, F., Ma, L., Saveanu, C., and Jacquier, A. (2015). Quality control of transcription start site selection by nonsense-mediated-mRNA decay. Elife 4. 10.7554/eLife.06722.

78. Lubas, M., Andersen, P.R., Schein, A., Dziembowski, A., Kudla, G., and Jensen, T.H. (2015). The human nuclear exosome targeting complex is loaded onto newly synthesized RNA to direct early ribonucleolysis. Cell Rep 10, 178–192. 10.1016/j.celrep.2014.12.026.

79. Gable, D.L., Gaysinskaya, V., Atik, C.C., Talbot, C.C., Jr., Kang, B., Stanley, S.E., Pugh, E.W., Amat-Codina, N., Schenk, K.M., Arcasoy, M.O., et al. (2019). ZCCHC8, the nuclear exosome targeting component, is mutated in familial pulmonary fibrosis and is required for telomerase RNA maturation. Genes Dev 33, 1381–1396. 10.1101/gad.326785.119.

80. Cordiner, R.A., Dou, Y., Thomsen, R., Bugai, A., Granneman, S., and Heick Jensen, T. (2023). Temporal-iCLIP captures co-transcriptional RNA-protein interactions. Nat Commun 14, 696. 10.1038/s41467-023-36345-y.

81. Wu, Y., Liu, W., Chen, J., Liu, S., Wang, M., Yang, L., Chen, C., Qi, M., Xu, Y., Qiao, Z., et al. (2019). Nuclear Exosome Targeting Complex Core Factor Zcchc8 Regulates the Degradation of LINE1 RNA in Early Embryos and Embryonic Stem Cells. Cell Rep 29, 2461–2472 e2466. 10.1016/j.celrep.2019.10.055.

82. Garland, W., Muller, I., Wu, M., Schmid, M., Imamura, K., Rib, L., Sandelin, A., Helin, K., and Jensen, T.H. (2022). Chromatin modifier HUSH co-operates with RNA decay factor NEXT to restrict transposable element expression. Mol Cell 82, 1691–1707 e1698. 10.1016/j.molcel.2022.03.004.

83. Falk, S., Finogenova, K., Melko, M., Benda, C., Lykke-Andersen, S., Jensen, T.H., and Conti, E. (2016). Structure of the RBM7-ZCCHC8 core of the NEXT complex reveals connections to splicing factors. Nat Commun 7, 13573. 10.1038/ncomms13573.

84. Imamura, K., Garland, W., Schmid, M., Jakobsen, L., Sato, K., Rouviere, J.O., Jakobsen, K.P., Burlacu, E., Lopez, M.L., Lykke-Andersen, S., et al. (in revision). A functional connection between the Microprocessor and a variant NEXT complex.

85. Trippe, R., Guschina, E., Hossbach, M., Urlaub, H., Luhrmann, R., and Benecke, B.J. (2006). Identification, cloning, and functional analysis of the human U6 snRNA-specific terminal uridylyl transferase. RNA 12, 1494–1504. 10.1261/rna.87706.

86. Trippe, R., Sandrock, B., and Benecke, B.J. (1998). A highly specific terminal uridylyl transferase modifies the 3’-end of U6 small nuclear RNA. Nucleic Acids Res 26, 3119–3126. 10.1093/nar/26.13.3119.

87. Trippe, R., Richly, H., and Benecke, B.J. (2003). Biochemical characterization of a U6 small nuclear RNA-specific terminal uridylyltransferase. Eur J Biochem 270, 971–980. 10.1046/j.1432-1033.2003.03466.x.

88. Heo, I., Ha, M., Lim, J., Yoon, M.J., Park, J.E., Kwon, S.C., Chang, H., and Kim, V.N. (2012). Mono-uridylation of pre-microRNA as a key step in the biogenesis of group II let-7 microRNAs. Cell 151, 521–532. 10.1016/j.cell.2012.09.022.

89. Faehnle, C.R., Walleshauser, J., and Joshua-Tor, L. (2017). Multi-domain utilization by TUT4 and TUT7 in control of let-7 biogenesis. Nat Struct Mol Biol 24, 658–665. 10.1038/nsmb.3428.

90. Kim, B., Ha, M., Loeff, L., Chang, H., Simanshu, D.K., Li, S., Fareh, M., Patel, D.J., Joo, C., and Kim, V.N. (2015). TUT7 controls the fate of precursor microRNAs by using three different uridylation mechanisms. EMBO J 34, 1801–1815. 10.15252/embj.201590931.

91. Hagan, J.P., Piskounova, E., and Gregory, R.I. (2009). Lin28 recruits the TUTase Zcchc11 to inhibit let-7 maturation in mouse embryonic stem cells. Nat Struct Mol Biol 16, 1021–1025. 10.1038/nsmb.1676.

92. Heo, I., Joo, C., Kim, Y.K., Ha, M., Yoon, M.J., Cho, J., Yeom, K.H., Han, J., and Kim, V.N. (2009). TUT4 in concert with Lin28 suppresses microRNA biogenesis through pre-microRNA uridylation. Cell 138, 696–708. 10.1016/j.cell.2009.08.002.

93. Zhang, E., Khanna, V., Dacheux, E., Namane, A., Doyen, A., Gomard, M., Turcotte, B., Jacquier, A., and Fromont-Racine, M. (2019). A specialised SKI complex assists the cytoplasmic RNA exosome in the absence of direct association with ribosomes. EMBO J 38, e100640. 10.15252/embj.2018100640.

94. Matera, A.G., and Wang, Z. (2014). A day in the life of the spliceosome. Nat Rev Mol Cell Biol 15, 108–121. 10.1038/nrm3742.

95. Neuenkirchen, N., Chari, A., and Fischer, U. (2008). Deciphering the assembly pathway of Sm-class U snRNPs. FEBS Lett 582, 1997–2003. 10.1016/j.febslet.2008.03.009.

96. Tudek, A., Lloret-Llinares, M., and Jensen, T.H. (2018). The multitasking polyA tail: nuclear RNA maturation, degradation and export. Philos Trans R Soc Lond B Biol Sci 373. 10.1098/rstb.2018.0169.

97. Stewart, M. (2019). Polyadenylation and nuclear export of mRNAs. J Biol Chem 294, 2977–2987. 10.1074/jbc.REV118.005594.

98. Passmore, L.A., and Coller, J. (2022). Roles of mRNA poly(A) tails in regulation of eukaryotic gene expression. Nat Rev Mol Cell Biol 23, 93–106. 10.1038/s41580-021-00417-y.

99. Huang, L., Li, G., Du, C., Jia, Y., Yang, J., Fan, W., Xu, Y.Z., Cheng, H., and Zhou, Y. (2023). The polyA tail facilitates splicing of last introns with weak 3’ splice sites via PABPN1. EMBO Rep 24, e57128. 10.15252/embr.202357128.

100. Bresson, S.M., and Conrad, N.K. (2013). The human nuclear poly(a)-binding protein promotes RNA hyperadenylation and decay. PLoS Genet 9, e1003893. 10.1371/journal.pgen.1003893.

101. Contreras, X., Depierre, D., Akkawi, C., Srbic, M., Helsmoortel, M., Nogaret, M., LeHars, M., Salifou, K., Heurteau, A., Cuvier, O., and Kiernan, R. (2023). PAPgamma associates with PAXT nuclear exosome to control the abundance of PROMPT ncRNAs. Nat Commun 14, 6745. 10.1038/s41467-023-42620-9.

102. Meola, N., and Jensen, T.H. (2017). Targeting the nuclear RNA exosome: Poly(A) binding proteins enter the stage. RNA Biol 14, 820–826. 10.1080/15476286.2017.1312227.

103. Silla, T., Karadoulama, E., Makosa, D., Lubas, M., and Jensen, T.H. (2018). The RNA Exosome Adaptor ZFC3H1 Functionally Competes with Nuclear Export Activity to Retain Target Transcripts. Cell Rep 23, 2199–2210. 10.1016/j.celrep.2018.04.061.

104. Shcherbik, N., Wang, M., Lapik, Y.R., Srivastava, L., and Pestov, D.G. (2010). Polyadenylation and degradation of incomplete RNA polymerase I transcripts in mammalian cells. EMBO Rep 11, 106–111. 10.1038/embor.2009.271.

105. Heng, J., Shi, B., Zhou, J.Y., Zhang, Y., Ma, D., Yang, Y.G., and Liu, F. (2023). Cpeb1b-mediated cytoplasmic polyadenylation of shha mRNA modulates zebrafish definitive hematopoiesis. Proc Natl Acad Sci U S A 120, e2212212120. 10.1073/pnas.2212212120.

106. Laishram, R.S. (2014). Poly(A) polymerase (PAP) diversity in gene expression--star-PAP vs canonical PAP. FEBS Lett 588, 2185–2197. 10.1016/j.febslet.2014.05.029.

107. Yamagishi, R., Tsusaka, T., Mitsunaga, H., Maehata, T., and Hoshino, S. (2016). The STAR protein QKI-7 recruits PAPD4 to regulate post-transcriptional polyadenylation of target mRNAs. Nucleic Acids Res 44, 2475–2490. 10.1093/nar/gkw118.

108. Barnard, D.C., Ryan, K., Manley, J.L., and Richter, J.D. (2004). Symplekin and xGLD-2 are required for CPEB-mediated cytoplasmic polyadenylation. Cell 119, 641–651. 10.1016/j.cell.2004.10.029.

109. Hojo, H., Yashiro, Y., Noda, Y., Ogami, K., Yamagishi, R., Okada, S., Hoshino, S.I., and Suzuki, T. (2020). The RNA-binding protein QKI-7 recruits the poly(A) polymerase GLD-2 for 3’ adenylation and selective stabilization of microRNA-122. J Biol Chem 295, 390–402. 10.1074/jbc.RA119.011617.

110. Millevoi, S., and Vagner, S. (2010). Molecular mechanisms of eukaryotic pre-mRNA 3’ end processing regulation. Nucleic Acids Res 38, 2757–2774. 10.1093/nar/gkp1176.

111. Mandel, C.R., Bai, Y., and Tong, L. (2008). Protein factors in pre-mRNA 3’-end processing. Cell Mol Life Sci 65, 1099–1122. 10.1007/s00018-007-7474-3.

112. Lee, E.S., Smith, H.W., Wolf, E.J., Guvenek, A., Wang, Y.E., Emili, A., Tian, B., and Palazzo, A.F. (2022). ZFC3H1 and U1-70K promote the nuclear retention of mRNAs with 5’ splice site motifs within nuclear speckles. RNA 28, 878–894. 10.1261/rna.079104.122.

113. Parker, R. (2012). RNA degradation in Saccharomyces cerevisae. Genetics 191, 671–702. 10.1534/genetics.111.137265.

114. Andersson, R., Refsing Andersen, P., Valen, E., Core, L.J., Bornholdt, J., Boyd, M., Heick Jensen, T., and Sandelin, A. (2014). Nuclear stability and transcriptional directionality separate functionally distinct RNA species. Nat Commun 5, 5336. 10.1038/ncomms6336.

115. Schwalb, B., Michel, M., Zacher, B., Fruhauf, K., Demel, C., Tresch, A., Gagneur, J., and Cramer, P. (2016). TT-seq maps the human transient transcriptome. Science 352, 1225–1228. 10.1126/science.aad9841.

