## Supplemental Tables for "RNA 3’end tailing safeguards cells against products of pervasive transcription termination"

**Table S1. siRNA sequences**

| siRNA target | sequence |
|---|---|
| DIS3L2 | GGUUGAUGGUGUUAAGAAAdTdT |
| NXF1 | UGAGCAUGAUUCAGAGCAAdTdT |
| PAPOLA | GAUUAGGAGUGCAUACAAAdTdT |
| PAPOLG | GGAGAAACAGAAAGGAAUAdTdT |
| PHAX | GAGUAUAUAGCACAGGAUUUAdTdT |
| SKIV2L | GGAGAUAGACUUUGAGAAAdTdT |
| ZCCHC8 | GGAAUGUACCUCAGGAUAAdTdT |
| ZFC3H1 | GAUUAGAGUCCAUGAUUAAdTdT |
| DIS3L | CGUAAAGACUUGAGGAAAAdTdT |
| Luc | CUUACGCUGAGUACUUCGAdTdT |
| DDX39A | AAAGGCCUAGCCAUCACUUUUdTdT |
| DDX39B | AAGGGCUUGGCUAUCACAUUUdTdT |
| siTENT4A | CUACGGUACCAAUAAUAAAdTdT |
| siTENT4B | GCGCUGACGUCCAGAUAUUdTdT |
| siTUT4 | GGAGAAACGACAUAAGAAAdTdT |
| siTUT7 | GAUAAGUAUUCGUGUCAAAdTdT |
| siTENT1 | GUGUGUUUGUCAGUGGCUUdTdT |
| siTENT2 | CAAUAUUGUUGGAAUAAGAdTdT |

**Table S2. Primary antibodies**

| Antibody | Source | Identifier | RRID |
|---|---|---|---|
| Puromycin | Merck Millipore | Cat#MABE343 | AB_2566826 |
| SKIV2L | Proteintech | Cat#11462-1-AP | AB_2187472 |
| DIS3L2 | Sigma-Aldrich | Cat#HPA035796 | AB_2674786 |
| Alpha-Tubulin | Rockland | Cat#600-401-880 | AB_2137000 |
| NXF1 | Abcam | Cat#ab129160 | AB_11142853 |
| DIS3L | Santa Cruz | Cat#sc-398739 | Not available |
| DDX39A/B | Santa Cruz | Cat#sc-271395 | AB_10609494 |
| TUT7 (ZCCHC6) | Sigma-Aldrich | Cat#HPA020600 | AB_1858985 |
| ZCCHC8 | Novus Biologicals | Cat#NB100-94995 | AB_1262274 |
| ZFC3H1 | Sigma-Aldrich | Cat#HPA-007151 | AB_1846133 |
| RRP40 | Santa Cruz | Cat#SC-101038 | AB_1122700 |
| Actin (ACTB) | Sigma-Aldrich | Cat#A2228 | AB_476697 |
| PAPOLG | Kindly provided by Georges Martin | | |
| PAPOLA | Kindly provided by Georges Martin | | |
| PHAX | Kindly provided by Edouard Bertrand | | |
| CBP80 | Kindly provided by Elisa Izaurralde | | |
| CBP20 | Kindly provided by Elisa Izaurralde | | |
| GFP | Kindly provided by John LaCava | | |

**Table S3. Primer sequences**

| **primers for qPCR** | |
|---|---|
| **Primer name** | **Sequence** |
| GAPDH F | GAGTCAACGGATTTGGTCGT |
| GAPDH R | TGAGGTGAATCAAGGGGTCA |
| proTTC32 F | GACCCGCCTTCCTCTTTTCC |
| proTTC32 R | TAAGAAACAGAACCGCGGCT |
| proSERINC3 F | TCTGCGAACTAAAGGATGAGTCC |
| proSERINC3 R | TGCACCCTTAGAGCCATTCA |
| proRBM39 F | CGGCCTCCCACACAGATG |
| proRBM39 R | TCTCCACCCTCAGCGGATAA |
| proMRPL1 F | ACCCTGTGTGTTTCGTGGTT |
| proMRPL1 R | AAAGTGGCGGAGGAAGCTAG |
| proHNRNPC F | GAAGCCTCCGTTGTGCAATG |
| proHNRNPC R | GAGAAAGGGTCTGGGGTTGG |
| proRHBDD2 F | CCAGCACCGTGTTTGATTCC |
| proRHBDD2 R | TGCAAGCTTCGTTTATTTTCGT |
| proMIPOL1 F | TCGAACTTAGTCTGCTGCGG |
| proMIPOL1 R | TCCGCGTTCTCCCTCTTCTA |
| proMRPS14 F | CCCTTCTCTCCCTACCCACA |
| proMRPS14 R | CTCTTGCCCTGTTCCTGGTC |
| TUT4 F | AAAAGGGACCCAGTTTACTGTTG |
| TUT4 R | GTCCGATACGTCTTCAATTCCTG |
| TENT1 F | TGTGTTTTTGTCGGTACAGGAG |
| TENT1 R | AGCTCGTAGTTCTACCAGGTG |
| TENT4A F | GCCTATATCGCCAAAGAGGAGA |

| | |
|---|---|
| TENT4A R | CTCCGGCCAACGTCATTCC |
| TENT4B F | GACATCGACCTAGTGGTGTTTG |
| TENT4B R | CGACTTTGTGTTTCCGAAGAGC |
| TENT2 F | ACCTACTGTTTATTCACACCAGC |
| TENT2 R | GCCGTTTACCGTCAAGAGGAA |
| Spike 1# F | AAGTTCGAAATCTAGGACGCA |
| Spike 1# R | GTACCCGGGAAAATCCTAG |
| Spike 2# F | AGACGCAGGCACAACTCTTT |
| Spike 2# R | CGCACGCCGAATGATGAAAC |
| **Primers for making RNA *in vitro* transcription templates** | |
| Spike 1# T7 F | TAATACGACTCACTATAGGGACTGACGAAGCTCAACATGCTG |
| Spike 1# R | GTACCCGGGAAAATCCTAG |
| Spike 1# 1U R | AGTACCCGGGAAAATCCTAG |
| Spike 1# 2U R | AAGTACCCGGGAAAATCCTAG |
| Spike 1# 3U R | AAAGTACCCGGGAAAATCCTAG |
| Spike 1# 4U R | AAAAGTACCCGGGAAAATCCTAG |
| Spike 1# 5U R | AAAAAGTACCCGGGAAAATCCTAG |
| Spike 2# T7 F | TAATACGACTCACTATAGGGTCAAAAAACTAGACGAATGGAG |
| Spike 2# R | CGCACGCCGAATGATGAAAC |
| **Primers for RT-PCR** | |
| $dT_{20}$-VN | TTTTTTTTTTTTTTTTTTTTVN |
| $dT_{20}$-$A_4$ | TTTTTTTTTTTTTTTTTTTTAAAA |
